# Sustained Generation of Neurons Destined for Neocortex with Oxidative Metabolic Upregulation upon Filamin Abrogation

**DOI:** 10.1101/2023.08.09.552667

**Authors:** Caroline A. Kopsidas, Clara C. Lowe, Dennis P. McDaniel, Xiaoming Zhou, Yuanyi Feng

## Abstract

Neurons in the neocortex are generated during embryonic development. While the adult ventricular-subventricular zone (V-SVZ) contains cells with neural stem/progenitors’ characteristics, it remains unclear whether it has the capacity of producing neocortical neurons. Here we show that the generation of neurons exhibiting transcriptomic resemblance to neurons of the upper cortical layers continues in the V-SVZ of mouse models of a human condition known as periventricular heterotopia by abrogating filamin. We found such surplus neurogenesis was associated with V-SVZ’s transcriptional upregulation of oxidative phosphorylation, mitochondrial biogenesis, and increased vascularization. Our spatial transcriptomics analysis also showed that the neurogenic activation of V-SVZ was coupled with enriched expression of genes in diverse pathways for energetics, signaling, neuronal activities, and metabolic turnovers of nucleic acids and proteins in upper cortical layers. These findings support the potential of generating neocortical neurons in adulthood through enhancing brain-wide vascular circulation, aerobic ATP synthesis, and neuronal vitality.

## Introduction

Neurons in the cerebral cortex are generated during embryogenesis following a precisely orchestrated developmental program. This program ensures both the timely production of cortical excitatory neurons from neural progenitor cells (NPCs) in the dorsal ventricular-subventricular zone (V-SVZ) situated adjacent to the brain ventricles and the subsequent migration of newborn neurons to form laminated neocortex underneath the brain surface(Gotz and Huttner, 2005; Noctor et al., 2004; Rakic, 1972; Taverna et al., 2014). With sufficient neurons being made to sustain cortical function throughout life, NPCs turn off neurogenesis and switch to making astrocytes and oligodendrocytes to support neuronal function(Alves et al., 2002; Qian et al., 2000; Toma and Hanashima, 2015). This NPC fate switch occurs before birth in both humans and rodents. Although cell proliferation persists in V-SVZ of juvenile and adult brains of humans, primates, and rodents(Kriegstein and Alvarez-Buylla, 2009; Mizrak et al., 2019; Sanai et al., 2004), evidence for neurogenic capacities in adulthood is only limited to producing olfactory neurons in rodents(Alvarez-Buylla and Garcia-Verdugo, 2002; Duque et al., 2022; Lim and Alvarez-Buylla, 2016; Nano and Bhaduri, 2022). As olfaction is less pivotal for sensory input in humans than rodents, it remains unclear whether cortical neurons can be made in the postnatal or adult V-SVZ to have an impact on human brain function and disorders. Thus, suitable experimental models and direct evidence are necessary to determine the propensity, mechanisms, and the functional implications of neurogenesis in the adult V-SVZ.

By studying the human brain developmental disorder periventricular heterotopia (PH) through murine models of Flna loss of function (LOF), we have shown previously that a “surplus” population of neurons can be made independent of embryonic neocortical neurogenesis and neuronal migration(Houlihan et al., 2016). PH, also known as periventricular nodular heterotopia or subependymal grey matter heterotopia, is a brain malformation characterized by abnormal clumping of neuronal nodules along the wall of lateral ventricles(Barkovich and Kjos, 1992; Barkovich and Kuzniecky, 2000; Eksioglu et al., 1996). Although many PH cases are sporadic, LOF mutations of the X-linked gene *FLNA* account for 20-30% of instances(Chen and Walsh, 1993; Fox et al., 1998; Lange et al., 2015; Parrini et al., 2006; Sheen et al., 2001). The affected individuals with *FLNA* mutations are mostly females; males in the pedigree may be affected but are often aborted spontaneously, presumably due to cardiovascular defects(Bernstein et al., 2011; Eksioglu *et al*., 1996; Reinstein et al., 2013). The most notable neurological symptom of PH is developing epilepsy beginning in mid-adolescence(Battaglia et al., 2006; d’Orsi et al., 2004; Huttenlocher et al., 1994; Lange *et al*., 2015). Although some affected individuals may present mild cognitive, behavioral, or reading deficits(Chang et al., 2005; Felker et al., 2011; Reinstein et al., 2012), the disease can often be asymptomatic or associated with little or no impairments on intellectual abilities(Chen and Walsh, 1993; Eisenbiegler and Brown, 2021; Lange *et al*., 2015; Lv et al., 2022; Parrini *et al*., 2006). In line with the minimal cognitive deficits, alterations in overall morphology and neocortical structure of the affected brains are also insignificant despite the presence of a large number of ectopic neurons (Ferland and Guerrini, 2009; Guerrini et al., 2004; Parrini et al., 2011; Poussaint et al., 2000; Reinstein *et al*., 2012), which argues that the generation of periventricular neurons has minimal impairments to the normal developmental program for cortical neurogenesis and neuronal migration.

*FLNA* encodes a 280kDa filamin A (FLNa) protein and is joined by *FLNB* and *FLNC* to form the filamin family(Nakamura et al., 2011; Wang and Singer, 1977; Weihing, 1985). While the expression of *FLNC* is specific to skeletal, cardiac, and smooth muscle, *FLNA* and *FLNB* are both expressed ubiquitously. The three filamin proteins share a similar structure including an N-terminal actin binding domain, two hinge domains, 24 immunoglobulin-like (Ig-like) repeats, and a C-terminal dimerization domain via the last repeat, allowing filamins to crosslink actin filaments as homo- or heterodimers(Gorlin et al., 1990; Sheen et al., 2002; Weihing, 1988). Besides actin, filamins have been reported to interact through the Ig-like repeats with over a hundred proteins with diverse functions in cytoplasm and the nucleus(Feng and Walsh, 2004; Lamsoul et al., 2020; Modarres and Mofradt, 2014; Stossel et al., 2001), suggesting additional roles of filamins besides merely stabilizing actin cytoskeleton necessary for cell motility and migration. PH is traditionally regarded as a neuronal migration disorder caused by mosaic *FLNA* expression in heterozygous females. However, this doesn’t explain cases caused by germline *FLNA* mutations in males(Cannaerts et al., 2018; Guerrini *et al*., 2004; Kasper et al., 2013; Lv *et al*., 2022; Oegema et al., 2013; Reinstein *et al*., 2013; Walsh et al., 2017). Although the migration defect of *FLNA* mutant neurons may be compensated by *FLNB,* our analyses of mouse PH models have shown that Flna and Flnb double null mutations in NPCs results in PH without affecting cortical neuronal migration (Houlihan *et al*., 2016). Instead, we showed that periventricular neurons were made extraneously due to altered V-SVZ micro-environment from epithelium mesenchymal transition (EMT) of polarized radial glial NPCs with filamin deficiency. The abnormal EMT was caused by compromised attenuation of IgF and Vegf signaling between NPCs and nascent vasculature in mid gestation of Flna-Flnb double deficient embryos, leading to feed-forward escalation of both angiogenesis and neurogenesis. The resulting periventricular neuronal nodules were first detectable at birth but continued to grow in size in postnatal development(Houlihan *et al*., 2016). This timeline of periventricular heterotopia formation in filamin mutant mice agrees with the corresponding human condition, in which neuronal nodules could be seen before birth but usually become evident along with the onset of epileptic seizures in the second decade of life. These together suggest strongly that PH is a condition in which a surplus of neurons is generated in the late-embryonic and postnatal V-SVZ.

Intriguingly, consistent with a vascular input in the initiation of PH shown by filamin deficient mice, *FLNA* mutations often cause defects in the cardiovascular system besides the brain(Bandaru et al., 2021; Chen and Walsh, 1993; Feng et al., 2006; Houlihan *et al*., 2016; Kyndt et al., 2007; Reinstein *et al*., 2013) and blood vessels are considered a major neurogenic niche in the V-SVZ of adult mammalian brains(Quaresima et al., 2022; Shen et al., 2008). Apart from supplying oxygen and nutrients for brain energy metabolism, cells of the vasculature can secrete VEGF and other factors to promote NPCs neurogenesis(Jin et al., 2002; Licht et al., 2016; Shen et al., 2004; Tavazoie et al., 2008). Thus, with a microenvironment of abundant blood vessels in the V-SVZ, filamin deficient NPCs may fail to turn off the production of neurons after the pre-programmed window of cortical neurogenesis and neuronal migration. As a result, neurons destined for the neocortex are made continuously by the V-SVZ neurons after birth, forming heterotopic nodules locally.

In this study, we set out to test this hypothesis by determining whether the generation of neocortical neurons persists in the postnatal and adult V-SVZ with mouse model of PH, what constitutes the molecular networks underlying sustained production of periventricular neurons, and how neurons in periventricular nodules are distinctive compared to neurons in the neocortex. Our results demonstrate that with combined increases in cerebral vascularization, oxidative energy metabolism, and vitalized cortical neuronal activities resulting from filamin deficiency, a large number of neurons with transcriptome resemblance of upper layer cortical neurons can be generated in the postnatal and adult V-SVZ but stay in situ to form heterotopic nodules. These results provide a conceptual possibility of boosting V-SVZ neurogenesis by targeting FLNA to enhance vascular circulation and energetics at the brain systems-level.

## Results

### Neurons in periventricular heterotopia increase continuously in juvenile and adult Fln^KO^ mice

Due to the embryonic lethality of Flna null mutation and the redundancy of Flna with Flnb, PH, defined by the presence of ectopic neurons (NeuN+) along the wall of lateral ventricles, was not observed in mice in which Flna was conditionally abrogated in cortical neural stem/progenitors (NSPCs) by Emx-Cre or nestin-Cre (referred to as Flna^cKO^ mice hereafter), but was found in all mice with compound Flna^cKO^ and homozygous Flnb mutations (Flna^f/yCre+^; Flnb^-/-^ males or Flna^f/f^ ^Cre+^; Flnb^-/-^females, referred to as Fln^KO^ or PH mice hereafter). About 20% of mice with compound Flna^cKO^ and heterozygous Flnb mutations (Flna^f/yCre+^; Flnb^+/-^ males or Flna^f/fCre+^; Flnb^+/-^ females) also presented bilateral PH with similar developmental and brain anatomical phenotype. In all cases, PH becomes detectable at perinatal ages (Houlihan *et al*., 2016). We further found that periventricular neuronal nodules in Fln^KO^ mice grew continuously not only in early neonatal ages but also in young adulthood. By following a cohort of Fln^KO^ mice and their littermate control mice (Flna^f/f^ ^Cre-^; Flnb^+/-^ or Flna^f/f^ ^Cre-^; Flnb^-/-^) from weaning to 10 months of age, we found that the brain weight of Fln^KO^ mice was significantly greater than that of control mice at ages of 4 months or older (Figure 1A, B). Histological and immonohistological (IH) analyses of serial coronal brain sections demonstrated that periventricular nodules in Fln^KO^ mice were reminiscent of those observed in human PH cases with *FLNA* mutations. They were heterogeneous in size and shape, and frequently grew together, forming larger masses or band-like structures. Although the diameter of some neuronal nodules could be comparable to the thickness of the neocortex, cerebral cortical size, structure, and neuronal lamination were overall indistinguishable between Fln^KO^ and control brains (Figure C, D). Therefore, increased brain weight in Fln^KO^ mice, from weaning to midlife, results from progressive addition of neurons to periventricular nodules. Our analysis also included a set of female mice with compound heterozygous Flna^cKO^ and homozygous Flnb mutations (Flna^f/wt^ ^Cre+^; Flnb^-/-^). Similar to females with heterozygous *FLNA* LOF in the human condition, all female mice with mosaic Flna NSPC conditional abrogation and Flnb null mutations exhibited bi-lateral periventricular neuronal nodules, while Flnb abrogation alone (Flna^f/f^ ^Cre-^; Flnb^-/-^) was not found to cause neuronal heterotopia or other brain structure changes (Figure S1A).

**Figure 1.**
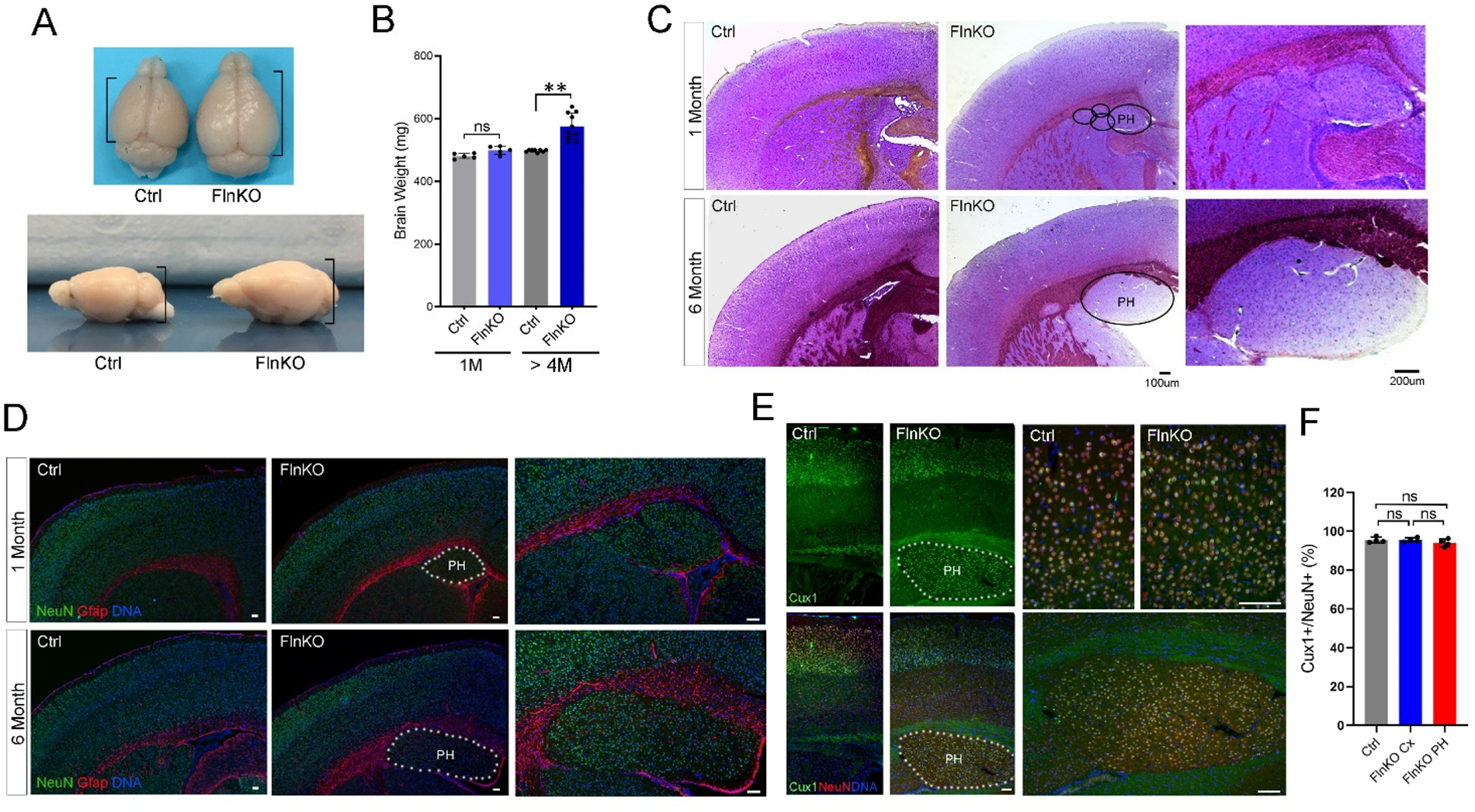
Periventricular Heterotopia continue to grow in Fln^KO^ mice after weaning. A. Representative brain images of a Fln^KO^ and a control mouse at 10 months of age. B. Brain weights of Fln^KO^ and control mice at ages of weaning (1M) and 4 month or older (≥ 4M), respectively. Shown are Mean ± SD. ** p < 0.01 by Student’s t-test. C. Representative images of Cresyl violet stained brain sections of a Fln^KO^ and control mice at weaning (1 month) and 6 months of age, respectively. The positions of PH are circled. D. Representative images of NeuN (green) and Gfap (red) double immunostained brain sections of Fln^KO^ and control mice at weaning (1 month) and 6 months of age, respectively. Nuclear DNA was stained with Hoechst 33342 and shown in blue. The positions of PH are indicated by dotted lines. E. Representative images of NeuN (red) and Cux1 (green) double immunostained brain sections of Fln^KO^ and control mice at 4 months of age. Nuclear DNA was stained with Hoechst 33342 and shown in blue. The positions of PH are indicated by dotted lines. F. Quantification (%) of Cux1+ neurons in the neocortex (Cx) of control mice, the neocortex of Fln^KO^ mice, and periventricular heterotopia (PH). Shown are Mean ± SD. Bars: 100 um or as indicated.

Starting at juvenile ages, the periventricular nodules in Fln^KO^ mice were predominantly composed of neurons surrounded by Gfap, an intermediate filament protein expressed by a subpopulation of astrocytes that are derived from embryonic NPCs and retain the potential of generating neuroblasts in the adult V-SVZ(Sanai *et al*., 2004; Tavazoie *et al*., 2008) (Figure 1D, Figure S1B, C). We also found that over 95% of periventricular neurons expressed Cux1, a marker for excitatory neurons of upper cortical layers that are generated towards the end of embryonic cortical neurogenesis (Figure 1E, F). This shared cell molecular identity between periventricular and late-born upper layer cortical neurons suggest that NSPCs in Fln^KO^ mice failed to shut down neurogenesis at the end of embryonic development and remained active in the postnatal and adult V-SVZ. The olfactory bulbs in Fln^KO^ mice were of normal size and structure, which further suggests that periventricular neurons were not migration-arrested olfactory neurons associated with the rodent-specific V-SVZ neurogenesis. Instead, they were likely cortical neurons produced by NSPCs that retained neurogenic propensity in the V-SVZ of postnatal and adult Fln^KO^ mice.

### Increased stemness and cellular activities in the V-SVZ of Fln^KO^ mice

To obtain further evidence for sustained generation of cortical neurons in adult Fln^KO^ mice, we examined the cellular composition and activities adjacent to periventricular neuronal nodules. Besides Gfap, we found that neurons in the nodular mass along the ventricular wall were surrounded by clusters of cells expressing the proliferation antigen Ki67 or the neuroblast marker Dcx (Figure 2A-C, Figure S2A). Although these markers for stemness and neurogenesis were also detectable in the V-SVZ of control mice, their abundance were substantially higher around periventricular neurons. Furthermore, we found cells in the V-SVZ of Fln^KO^ mice proliferate much more actively than those in control mice as evidenced by substantial increases in EdU incorporation (Figure 2D, E, Figure S2B). Notably, EdU+ cells in Fln^KO^ V-SVZ distributed around periventricular neurons along with Gfap and Dcx, which suggests that proliferative and neurogenic activities of Fln^KO^ NSPC were beyond the routine olfactory neurogenesis and largely dedicated towards generating periventricular neurons. Along with this notion, cells expressing Olig2+ and Sox10+, which mark oligodendrocytes and precursors, were also significantly elevated in the V-SVZ of Fln^KO^ compared to that of control mice (Figure 2F-I, Figure S2C, D). This complementary increase in periventricular neurogenesis and oligodendrocyte differentiation in the V-SVZ of Fln^KO^ mice indicates a coordinated elevation in stemness and cellular activities towards generating both neurons and their supporting cells.

**Figure 2.**
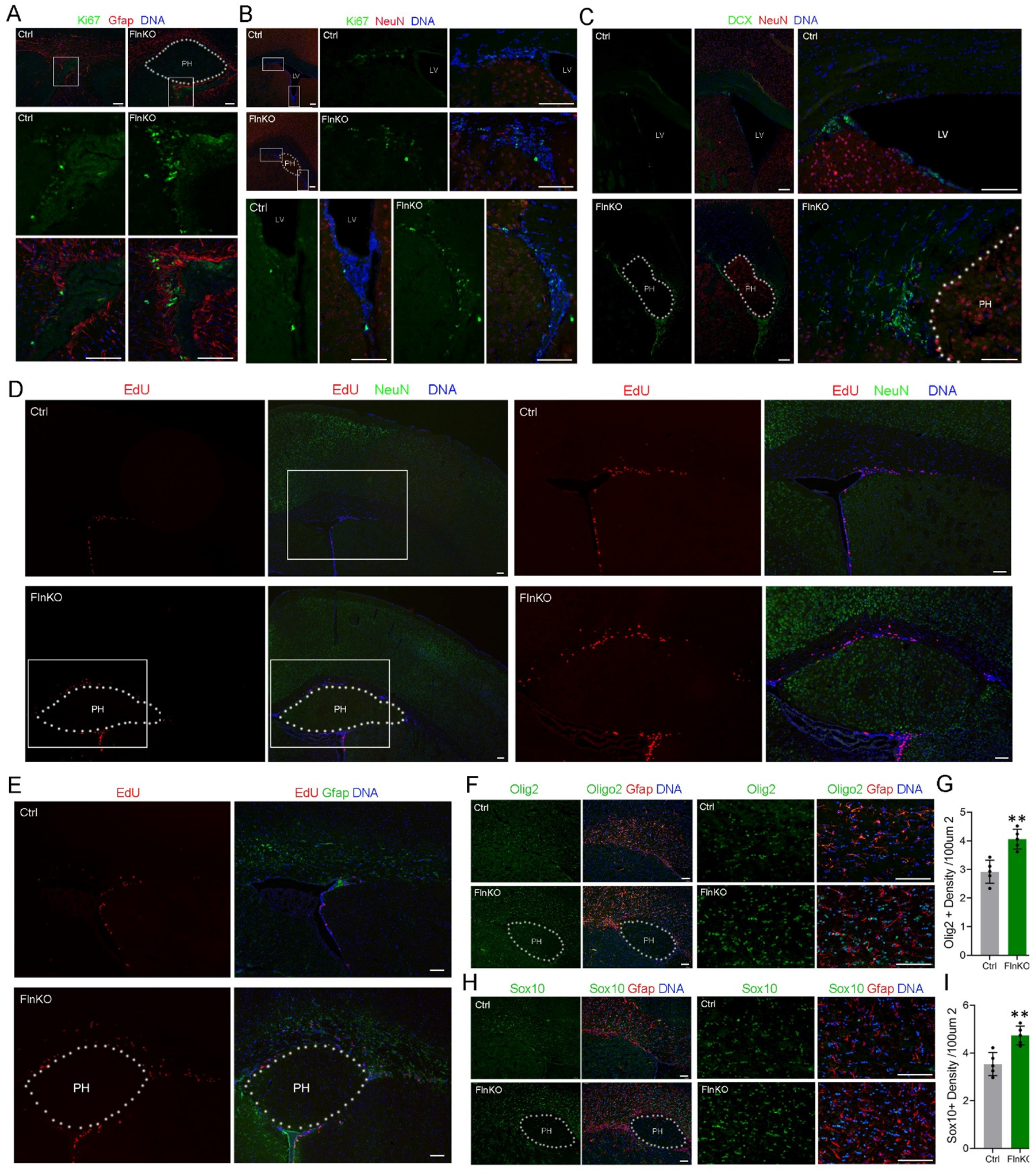
Increased stemness and cellular activity towards neurogenesis and glial differentiation in the V-SVZ of Fln^KO^ mice. A. Representative images of Ki67 (green) and Gfap (red) double immunostained brain sections of Fln^KO^ or control mice at 4 months of age. Nuclear DNA was stained with Hoechst 33342 and shown in blue. The position of PH is marked by dotted lines. B. Representative images of Ki67 (green) and NeuN (red) double immunostained brain sections of Fln^KO^ or control mice at 4 months of age. Nuclear DNA was stained with Hoechst 33342 and shown in blue. The position of PH is marked by dotted lines. C. Representative images of DCX (green) and NeuN (red) double immunostained brain sections of Fln^KO^ or control mice at 3 months of age. Nuclear DNA was stained with Hoechst 33342 and shown in blue. The position of PH is marked by dotted lines. D. Representative images of brain sections stained by EdU (red) and immunostained by anti-NeuN (green) of Fln^KO^ or control mice at 3 months of age. Nuclear DNA was stained with Hoechst 33342 and shown in blue. The position of PH is marked by dotted lines. E. Representative images of brain sections stained by EdU (red) stained and immunostained by anti-Gfap (green) of Fln^KO^ or control mice at 3 months of age. Nuclear DNA was stained with Hoechst 33342 and shown in blue. The position of PH is marked by dotted lines. F. Representative images of Olig2 (green) and Gfap (red) double immunostained brain sections of Fln^KO^ or control mice at 3 months of age. Nuclear DNA was stained with Hoechst 33342 and shown in blue. The position of PH is marked by dotted lines. G. Quantification of the density of Olig2+ cells in the dorsal V-SVZ of control or Fln^KO^ mice. Shown are Mean ± SD. ** p< 0.01 by Student’s t-test. H. Representative images of Sox10 (green) and Gfap (red) double immunostained brain sections of Fln^KO^ or control mice at 3 months of age. Nuclear DNA was stained with Hoechst 33342 and shown in blue. The position of PH is marked by dotted lines. I. Quantification of the density of Olig2+ cells in the dorsal V-SVZ of control or Fln^KO^ mice. Shown are Mean ± SD. ** p< 0.01 by Student’s t-test. Bars: 100 um or as indicated.

### Upregulation of genes mediating oxidative metabolism in the V-SVZ of Fln^KO^ mice

To determine the molecular basis and functional gene networks associated with periventricular neurogenesis at the brain systems level, we employed spatial transcriptomics to capture brain regional specific gene expression profiles. We prepared brain sections from Fln^KO^ and control mice at 3 months of age (n=3 each) and double stained them with fluorescent-conjugated antibodies to Gfap and NeuN. Guided by Gfap signals along the lateral ventricles, we selected 16 and 15 regions of interest (ROIs) from the dorsal V-SVZ of control and Fln^KO^ brains, respectively. Based on NeuN signals, we also selected 16 and 15 ROIs from upper cortical layers of Control and Fln^KO^ brains, respectively. In addition, Gfap-NeuN double staining allowed the selection of 14 ROIs from the periventricular nodules of Fln^KO^ brains (Figure S3A). We used the NanoString GeoMx DSP platform, by which RNA sequencing yielded expression data for 19,962 genes across all 76 sampled ROIs. Comparing regionally-specific ROIs between Fln^KO^ and control mice revealed 4,003 and 7,411 differentially expressed genes (DEGs) in the V-SVZ and upper cortex, respectively. In addition, we identified 612 DEGs between ROIs in periventricular nodules and upper cortical layers of Fln^KO^ mice (Figure 3A, Figure S3B-E). Therefore, despite the lack of discernible change in neocortical morphology, generation of periventricular neurons in Fln^KO^ mice was associated with remarkable transcriptome alterations in both V-SVZ and upper cortical layers.

**Figure 3.**
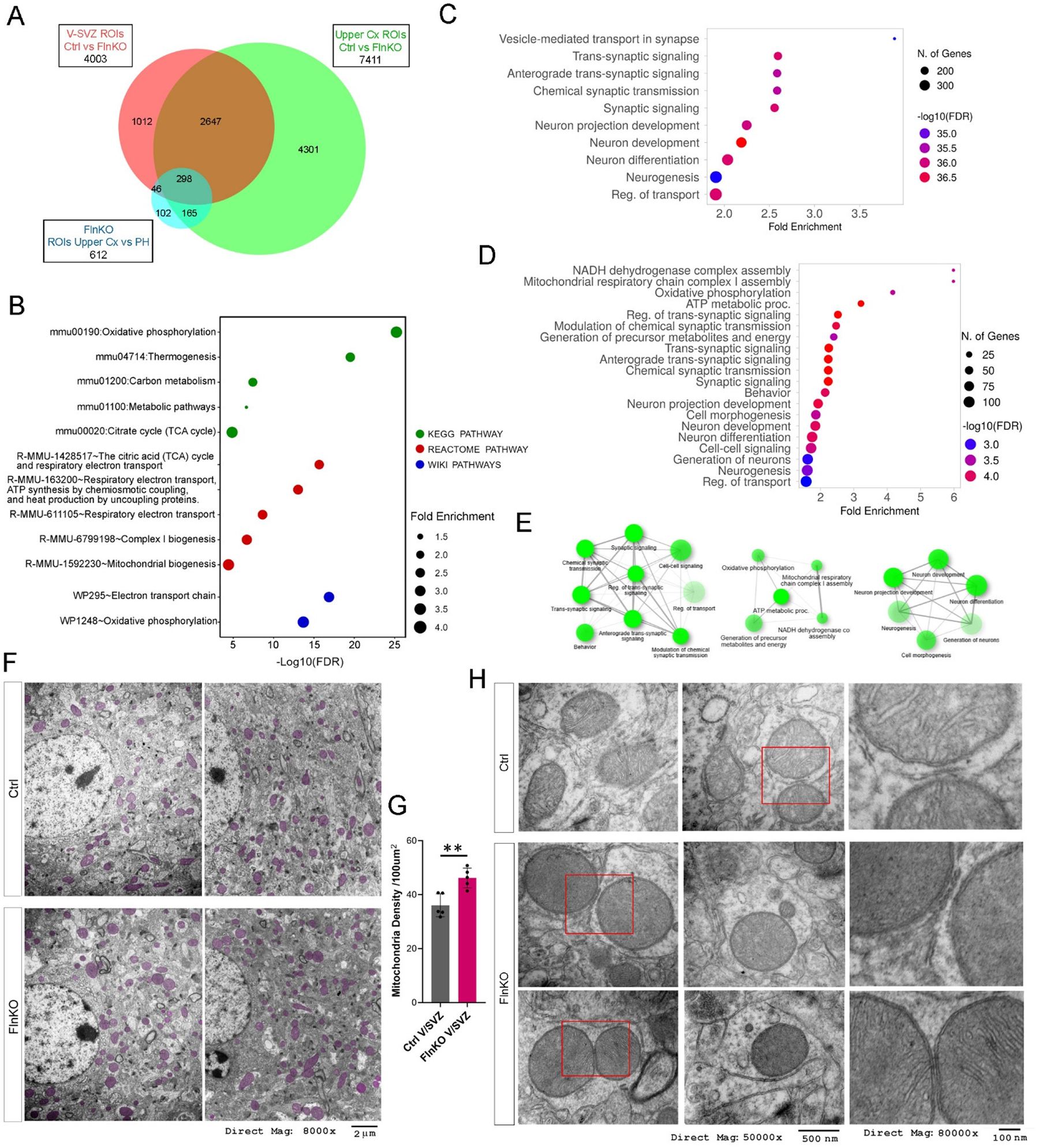
Increased V-SVZ neurogenesis in Fln^KO^ mice is associated with upregulation of oxidative energy generation and mitochondrial biogenesis. A. Venn diagram of differentially expressed genes (DEGs) identified by GeoMx DSP analysis of mouse whole transcriptome. A total of 76 regions of interests (ROIs) representing V-SVZ, upper cortical layers, and periventricular heterotopia were analyzed in Fln^KO^ and control mice at 3-4 months of age (biological replicate =3 each). Shown are numbers of DEGs from V-SVZ ROIs between Fln^KO^ (n=15) and control (n=16) mice, from upper cortical ROIs between Fln^KO^ (n=15) and control (n=16), and from periventricular heterotopia nodule ROIs (n=14) and upper cortical layer ROIs (n=15) of Fln^KO^ mice. B. Summary of KEGG, Reactome, and Wiki pathway enrichment analyses of upregulated DEGs across V-SVZ ROIs of Fln^KO^ mice compared to those of control mice. C. Gene Ontology Biological Processes (BP) terms enriched in upregulated DEGs across V-SVZ ROIs of Fln^KO^ mice compared to those of control mice. D. Gene Ontology Biological Processes (BP) terms enriched in DEGs exclusively upregulated across V-SVZ ROIs of Fln^KO^ mice compared to those of control mice. E. Network plots show the relationship of enriched BP terms shown in D. F. Representative electron micrographs of V-SVZ of Fln^KO^ and control mice at 5 months of age. Mitochondria are highlighted in mauve. G. Quantification of mitochondrial density in the soma of V-SVZ cells of Fln^KO^ and control mice at 5 months of age. Shown are Mean ± SD. ** p< 0.01 by Student’s t-test. H. Representative electron micrographs of mitochondria in V-SVZ of Fln^KO^ and control mice at 5 months of age. Images at higher magnification of boxed areas are included.

To assess the molecular profile underling neurogenic activation in the V-SVZ of Fln^KO^ mice, we first queried DEGs identified across V-SVZ ROIs between Fln^KO^ and control mice. Out of the total 4,003 DEGs, the 1,489 downregulated DEGs showed no significant enrichment in pathway analysis. In contrast, the 2,514 upregulated DEGs in Fln^KO^ V-SVZ were found enriched in many functional pathways and biological processes, among which the KEGG pathway of oxidative phosphorylation (OXPHOS) was most significantly overrepresented (FDR =1.2E-25; Fold Enrichment = 4.4) (Figure 3B, C, Figure S4). Remarkably, 65 of 133 genes participating in OXPHOS were upregulated in Fln^KO^ compared to control V-SVZ ROIs (Figure S4A), suggesting strongly that the increased V-SVZ stemness and neurogenic activities in Fln^KO^ mice were supported by elevating mitochondrial electron transport chain (ETC) function for aerobic energetics.

As 1,735 of DEGs upregulated across V-SVZ ROIs also showed significantly higher expression in upper cortical ROIs of Fln^KO^ mice (Figure 3A, Figure S3E), we assessed the 779 genes that were exclusively upregulated in the Fln^KO^ V-SVZ to determine the V-SVZ-specific impact. Our results of pathway and gene ontology (GO) enrichments analyses showed that these V-SVZ-specific DEGs represented those functional pathways and biological processes related to not only mitochondrial biogenesis and oxidative metabolic processes for ATP production but also neural developmental processes, such as generation, differentiation, and morphogenesis of neurons, or networking activities of synaptic signaling, transmission and behaviors (Figure 3D, E). The co-upregulation of genes for mitochondrial function and neural development in the Fln^KO^ V-SVZ suggests that the switch of NSPCs from quiescence to neurogenesis is coupled with a metabolic transition towards a state dominated by OXPHOS.

We next performed transmission electron microscopic analyses to reveal the mitochondrial ultrastructure underlying the metabolic transition associated with V-SVZ neurogenesis. We found that despite heterogeneity in size, morphology, and distribution, mitochondria in the V-SVZ of Fln^KO^ mice were not only significantly higher in density compared to those in the V-SVZ of control mice but also showed distinctive structural features (Figure 3F-H). These mitochondria frequently displayed a spherical shape containing densely packed cristae (Figure 3H). While the spherical morphology is a characteristic of mitochondria in stem cells, condensed mitochondrial cristae indicate strong ETC activity for ATP synthesis(Mannella, 2006; Naik et al., 2019). These data suggest that generation and functional maturation of new neurons in the V-SVZ of Fln^KO^ mice rely on a metabolic upgrade from anaerobic glycolysis to OXPHOS through mitochondrial biogenesis. As ATP is highly indispensable for neurons in the adult brain, our data imply that in order to fulfill the energetics demand of adult neurons, upregulating the efficiency of ATP synthesis is essential for committing NPCs to neuronal differentiation.

### Increased vascularization in Fln^KO^ brains supports system-wide vitalization of neuronal activity and plasticity

The energy substrates of the brain are solely supplied by vascular circulation which constantly delivers glucose and oxygen to permit high efficiency ATP synthesis by OXPHOS. Thus, upregulation of energetics in adult Fln^KO^ V-SVZ might be supported by angiogenesis as observed in the developing Fln^KO^ brain(Houlihan *et al*., 2016). Indeed, we found that the abundance of cerebral blood vessels was significantly higher in Fln^KO^ than in control mice (Figure 4A-C). Besides an overall increase in density, blood vessels in Fln^KO^ brains, especially those surrounding and inside the periventricular nodules, frequently showed substantially enlarged caliber (Figure 4A B, D). The enlarged vessels near heterotopic neurons were also found to be associated with increased propensity of EdU incorporation (Figure 4E), which supports the vascular involvement in local cellular proliferation, metabolism, communication, and angiogenesis. These results are fully in line with blood vessels being an essential constituent of the V-SVZ neurogenic niche. They suggest that increased vascularization in Fln^KO^ brains not only facilitates the delivery of substrates for OXPHOS but also increases the flux of numerous extrinsic factors, such as growth factors, hormones, and endothelium derived molecules, which are together necessary for NSPC self-renewal and neurogenic activation.

**Figure 4.**
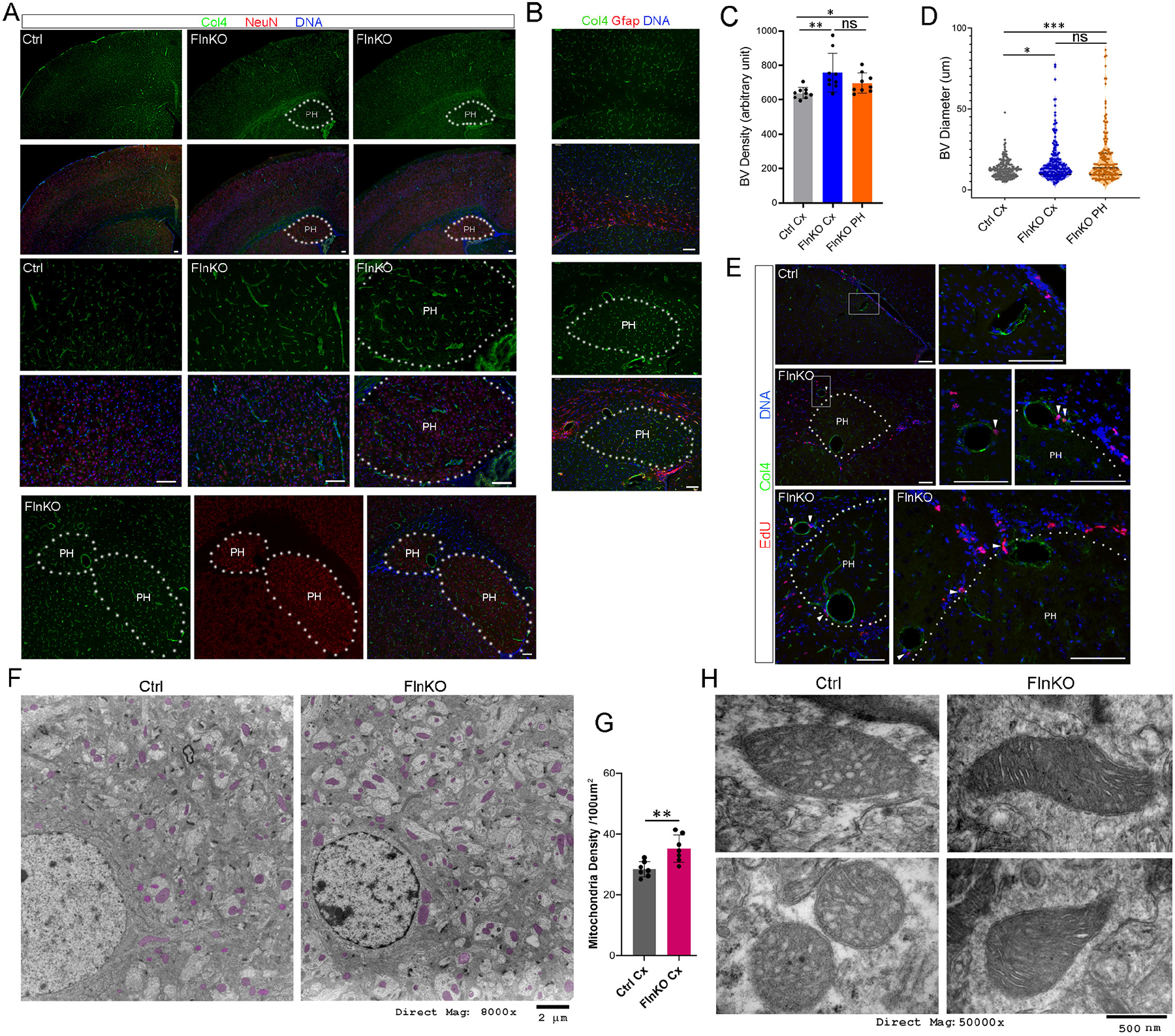
Increased vascularization and upregulation of energy metabolic activities throughout the neocortex of Fln^KO^ mice. A. Representative images of Collagen IV (Col4, green) and NeuN (red) double immunostained brain sections of Fln^KO^ or control mice at 3 months of age. Nuclear DNA was stained with Hoechst 33342 and shown in blue. The positions of PH are marked by dotted lines. B. Representative images of Collagen IV (Col4, green) and Gfap (red) double immunostained brain sections of Fln^KO^ or control mice at 4 months of age. Nuclear DNA was stained with Hoechst 33342 and shown in blue. The positions of PH are marked by dotted lines. C. Quantification of the density of blood vessels in neocortex (Cx) of control mice, neocortex of Fln^KO^ mice, and periventricular heterotopia (PH). Samples were from mice at 3-4 months of age (n ≥ 4 each genotype). Shown are Mean ± SD. *p< 0.05; ** p< 0.01 by Student’s t-test. D. Quantification of the caliber of blood vessels in neocortex (Cx) of control mice, neocortex of Fln^KO^ mice, and periventricular heterotopia (PH). Samples were from mice at 3-4 months of age (n ≥ 4 each genotype). Shown are violin plots with data points, median, and quartiles. *p< 0.05; *** p< 0.001 by Kolmogorov-Smirnov test. E. Representative images of brain sections of control and Fln^KO^ mice stained by EdU (red) stained and immunostained by anti-Col4 (green). Nuclear DNA was stained with Hoechst 33342 and shown in blue. The position of PH is marked by dotted lines. Arrowheads indicate EdU+ cells associated with enlarged blood vessels. F. Representative electron micrographs of the neocortex of Fln^KO^ and control mice at 5 months of age. Mitochondria are highlighted in mauve. G. Quantification of mitochondrial density in the neocortex of Fln^KO^ and control mice at 5 months of age. Shown are Mean ± SD. ** p< 0.01 by Student’s t-test. H. Representative electron micrographs of mitochondria in the neocortex of Fln^KO^ and control mice at 5 months of age. Bars: 100um or as indicated.

The increase in vascular abundance is expected to have an impact at the systems-level, affecting global brain metabolism and function. Consistent with this, our ultrastructure analysis revealed a significant increase in mitochondrial density in the neocortex of Fln^KO^ mice compared to that of control mice (Figure 4F, G). Similar to observed in the V-SVZ, mitochondria in the neocortex of Fln^KO^ brains showed densely packed cristae (Figure 4H), suggesting a more active state for aerobic ATP synthesis.

Metabolic activation in the neocortex of Fln^KO^ brain is also demonstrated by differential transcriptome profiles identified across ROIs of upper cortical layers between Fln^KO^ and control mice. Out of the total 7,411 DEGs, the 4,609 upregulated genes in upper cortex of Fln^KO^ mice showed significant over-representation of numerous pathways for metabolism, neuronal activity, signal transduction, and cellular structure regulations (Figure 3A, Figure S3, Figure 5A). These include not only OXPHOS but also KEGG and Reactome pathways for metabolic regulations of thermogenesis, the TCA cycle and respiratory chain transport, metabolism of RNA, metabolism of proteins, mRNA splicing, protein translation, signaling cascades of G-protein, cAMP, cGMP, AMPK, RAF/MAPK, ErbB, insulin, Wnt, oxytocin, Apelin, FoxO, estrogen, as well as neuronal activity pathways in long-term potentiation (LTP), long-term depression (LTD), glutamatergic synapse, dopaminergic synapse, GABAergic synapse, and cholinergic synapse activation (Figure 5A, Figure S5A-D).

**Figure 5.**
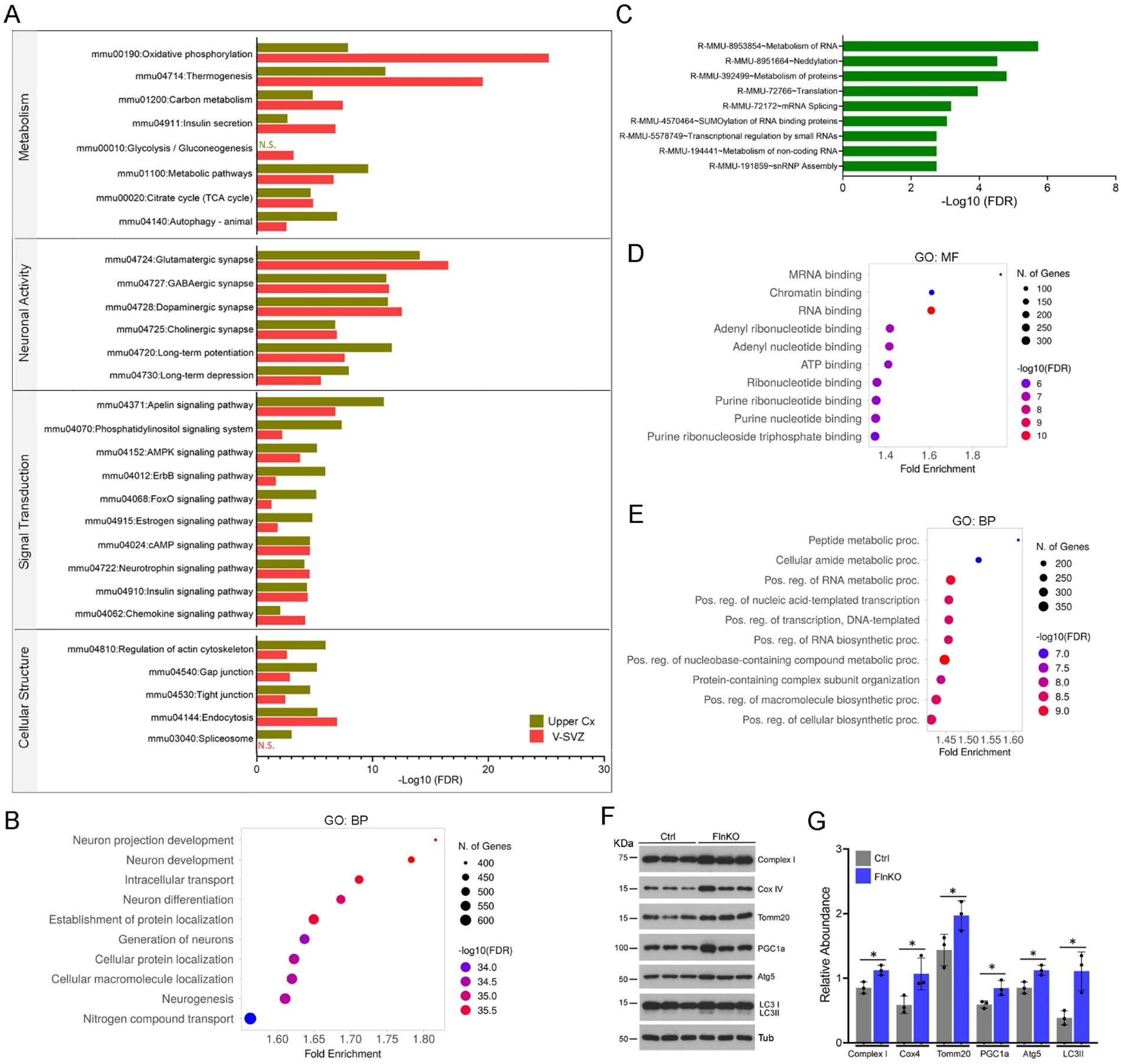
Transcriptional upregulation of genes mediating brain metabolic and neural activities in Fln^KO^ mice. A. KEGG pathway analysis of upregulated genes in V-SVZ ROIs and upper cortical ROIs of Fln^KO^ mice, respectively. Note the more significant upregulation of genes mediating OXPHOS and thermogenesis in V-SVZ ROIs than upper cortical ROIs, and the more significant upregulation of genes essential for autophagy, LTP, LTD, Apelin signaling, and spliceosome in upper cortical ROIs than V-SVZ ROIs in Fln^KO^ mice. B. Gene Ontology Biological Processes (BP) terms enriched in all DEGs upregulated across upper cortical ROIs of Fln^KO^ mice compared to those of control mice. C. Reactome pathways over-represented by DEGs exclusively upregulated across upper cortical ROIs of Fln^KO^ mice. D. Gene Ontology Molecular Function (MF) terms enriched in DEGs exclusively upregulated across upper cortical ROIs of Fln^KO^ mice. E. Gene Ontology Biological Processes (BP) terms enriched in DEGs exclusively upregulated across upper cortical ROIs of Fln^KO^ mice. F, G. Immunoblotting analysis of total cortical protein extracts. Shown are representative IB images and quantifications (Mean ± SD). * p< 0.05 by Student’s t-test.

As there is a substantial overlap in pathways over-represented by upregulated DEGs in V-SVZ and upper cortical layers of Fln^KO^ mice, we determined their brain regional-specificity. Although genes in pathways of OXPHOS, thermogenesis, insulin secretion, and gluconeogenesis were significantly upregulated in both V-SVZ and upper cortex of Fln^KO^ mice, their enrichments in V-SVZ were more pronounced than in upper cortical layers (Figure 5A), suggesting a stronger demand of upregulating aerobic glucose catabolism for V-SVZ neurogenesis. In contrast, genes mediating LTP, LTD, autophagy, and Apelin signaling were more significantly enriched in upper cortical than V-SVZ ROIs of Fln^KO^ mice (Figure 5A). LTP and LTD are important for synaptic communication and plasticity; autophagy is also crucial for neuronal activity-mediated metabolic recycling that may facilitate neuronal connectivity and remodeling; while the major role of Apelin signaling is promoting angiogenesis and vasodilation. Therefore, the co-upregulation of molecules in these pathways, along with increased cerebral vascularization in Fln^KO^ cortices, has a positive impact on synaptic activity, plasticity, and molecular turnover in cortical neurons. In line with this, biological processes over-represented by upper cortical ROIs in Fln^KO^ mice were predominantly related to nervous system development, including GO BP terms in neuron development, neuron differentiation, and intracellular transport (Figure 5B). Taken together, these data suggest that neurogenic activation in the V-SVZ in Fln^KO^ mice is coordinated with brain-wide boost of neuronal signaling and network activities through enhancing vascular circulation and function.

To further validate elevations in activity, plasticity, and metabolic turnover of Fln^KO^ cortical neurons, we queried the 2,874 DEGs that were exclusively increased in upper cortical ROIs (Figure S3E). We found transcripts specifically enriched in the upper cortex of Fln^KO^ mice over-represented those Reactome pathways largely associated with RNA and protein metabolic synthesis, modification, and turnover (Figure 5C). Likewise, GO enrichment analyses showed that these DEGs exclusively upregulated in Fln^KO^ upper cortices were significantly associated with MF and BP terms in RNA or nucleotide binding as well as RNA and peptide metabolic processes (Figure 5D, E). These metabolic activities are not only necessary for cell proliferation, differentiation, survival, and function, but are also essential for rapid turnover of synaptic proteins. Collectively, the transcriptome profiling data suggest that Fln^KO^ brains are vitalized not only with greater propensity of generating new neurons in the postnatal and adult V-SVZ but also with higher activity and plasticity of those cortical excitatory neurons born before birth.

Consistent with the role of filamin in mediating numerous protein-protein interactions encompassing the actin cytoskeleton and diverse cytoplasmic or nuclear molecules, upregulated genes in both upper cortical and V-SVZ ROIs of Fln^KO^ mice were associated with pathways regulating actin cytoskeleton, gap junction, tight junction, endocytosis, membrane trafficking, and protein-protein interactions at synapses (Figure 5A, Figure S5). In addition, GO analysis showed that upregulated DEGs in Fln^KO^ cortices were enriched in those CC terms essential for neurons, including synapses, dendrites, axons, myelin sheath, and mitochondrion, and those MP terms involving diverse molecular bindings of RNAs, proteins and protein complexes, protein kinases, transcriptional factors, chromatin, cytoskeleton, and nucleotides (Figure S5E, F). In contrast to the extensive functional annotation of upregulated genes, the 2,802 down-regulated DEGs across upper cortical ROIs of Fln^KO^ mice showed limited association with biological processes and functional pathways. Similar to the 1,489 down-regulated genes in V-SVZ ROIs, genes with significantly decreased expression in upper cortical layers of Fln^KO^ mice were only enriched strongly in biological processes mediating pheromone responses (Figure S5G, H).

Further corroborated with ultrastructural and transcriptome data, our immunoblotting (IB) analyses showed increased abundances of molecules for mitochondrial biogenesis and ETC in Fln^KO^ cortices (Figure 5F, G). These include molecules of mitochondrial respiratory chain complexes, mitochondrial outer membrane protein Tomm20, and PGC1α, the master regulator for mitochondria biogenesis. In correlation with autophagy elevation indicated by transcriptomics analysis, we found both Atg5 and LC3II protein were increased significantly in Fln^KO^ cortical tissues. Altogether, with increased vascularization, OXPHOS, and metabolic turnovers of RNA and proteins, both V-SVZ and cerebral cortex of adult Fln^KO^ mice appear in a relatively immature and hyperactive state, retaining more features of a youthful developing brain.

### Periventricular neurons show transcriptome resemblance with upper cortical neurons but reduced expression of neuronal activity genes

To determine the characteristics of neurons generated in the postnatal and adult V-SVZ, we asked how the neurons in periventricular nodules differ from upper cortical layers in Fln^KO^ mice, even though both populations are predominantly Cux1+ neurons (Figure 1E). We first compared their transcriptomic profiles to assess their molecular identity and activity. In contrast to the remarkable transcriptomic difference between upper cortical layers of Fln^KO^ and control mice, only 612 genes showed differential expression between upper cortical and periventricular ROIs in Fln^KO^ mice (Figure 3A). Therefore, upper cortical neurons in Fln^KO^ mice share a greater degree of molecular resemblance with periventricular neurons than with their upper cortical counterparts in control mice. Furthermore, the 612 DEGs did not include genes specific for olfactory neurons, which ruled out the possibility of excessive olfactory neurogenesis or failed migration of olfactory neurons in Fln^KO^ mice. Instead, we found that the 167 DEGs upregulated in periventricular nodules were moderately associated with morphogenesis, gliogenesis, ECM remodeling, and cell stimuli responses, while the 445 DEGs downregulated in periventricular nodules were significantly enriched in pathways and biological processes in neurodevelopment and synaptic activities, including synaptic signaling, neuronal system, neuron projection development, regulation of membrane potential, organelle localization, membrane trafficking, behavior, as well as Ca2+ regulation and signaling by Rho GTPase (Figure 6A, B, Figure S6A). These data indicated that the major difference between neurons in periventricular nodules and upper cortical layers lies in the functional activity instead of neuronal subtype identity. They suggest that although the postnatal and adult V-SVZ of Fln^KO^ mice retains the potential of generating neurons destined for the neocortex, neurons produced outside of the normal window of cortical development fail to become fully active as they are unable to integrate into the functional cortical circuitry.

**Figure 6.**
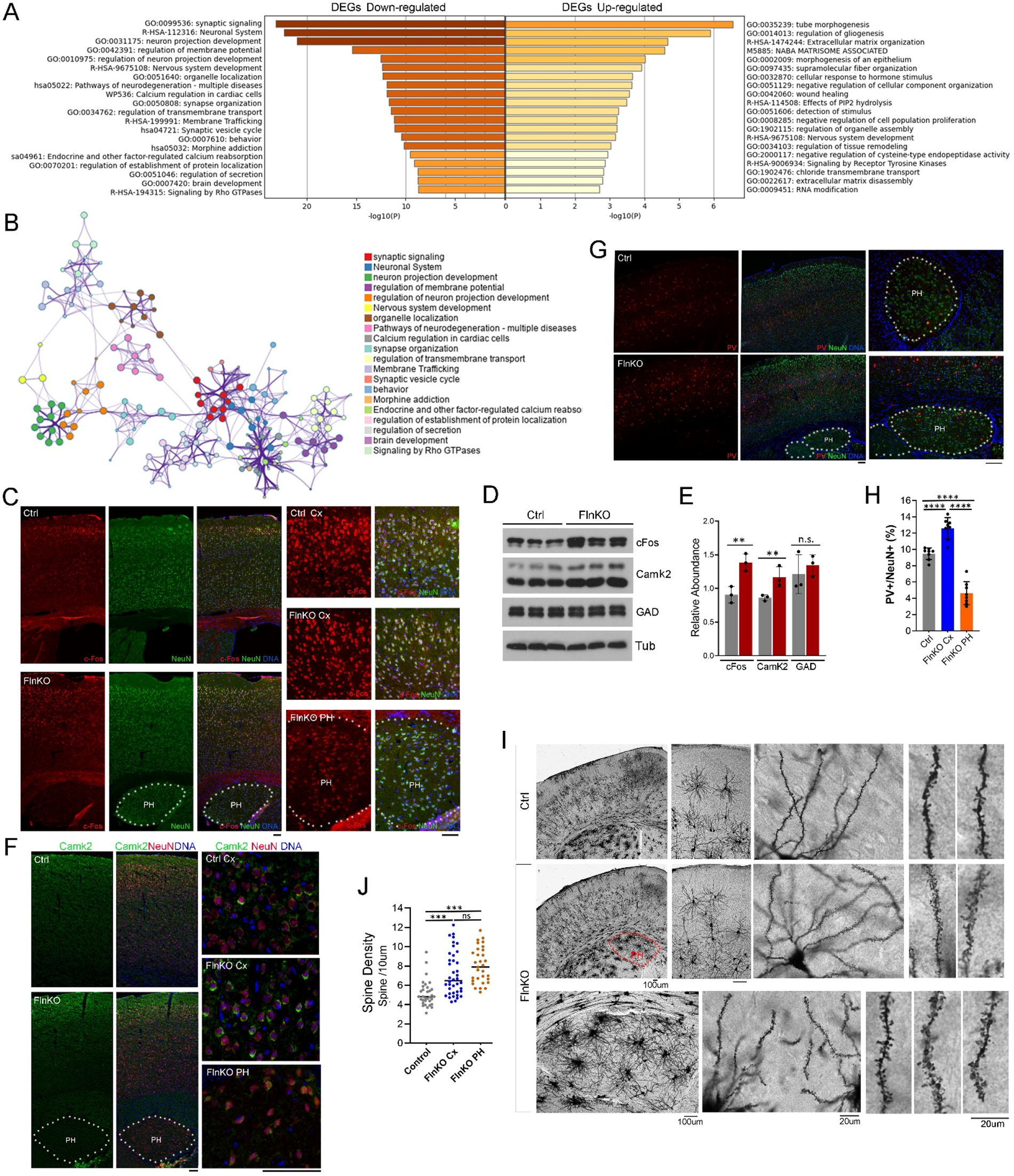
Periventricular neurons present a transcriptome profile overlapping with that of upper cortical neurons in Fln^KO^ mice but show decreased activity. A. Statistically enriched GO terms, KEGG or canonical pathways, and hallmark gene sets in genes upregulated and downregulated, respectively, in ROIs of periventricular nodules compared to ROIs of upper cortical layers of Fln^KO^ mice. B. Network of genes downregulated in ROIs across periventricular nodules compared to ROIs across upper cortical layers of Fln^KO^ mice. C. Representative images of cFos (red) and NeuN (green) double immunostained brain sections of Fln^KO^ or control mice at 3 months of age. Nuclear DNA was stained with Hoechst 33342 and shown in blue. Note the low cFos immunosignals in periventricular neurons. The position of PH is marked by dotted lines. D, E. Immunoblotting analysis of total cortical protein extracts. Shown are representative IB images and quantifications (Mean ± SD). ** p< 0.01 by Student’s t-test. F. Representative images of CamK2 (green) and NeuN (red) double immunostained brain sections of Fln^KO^ or control mice at 3 months of age. Nuclear DNA was stained with Hoechst 33342 and shown in blue. Note the low CamK2 immunosignals in periventricular neurons. The position of PH is marked by dotted lines. G. Representative images of Parvalbumin (PV, red) and NeuN (green) double immunostained brain sections of Fln^KO^ or control mice at 4 months of age. Nuclear DNA was stained with Hoechst 33342 and shown in blue. The position of PH is marked by dotted lines. H. Quantification of PV+ neurons (% out of total NeuN+) in the neocortex (Cx) of control mice, the neocortex of Fln^KO^ mice, and periventricular heterotopia (PH). Shown are Mean ± SD. **** p <0.0001 by Student’s t-test. I. Representative images of Golgi Cox stained Fln^KO^ and control brains for neuronal morphology and dendritic spine analyses. Note that dendritic spines in Fln^KO^ mice show relatively thin neck and fridge shape. J. Quantification of dendritic spine densities in the neocortex (Cx) of control mice, the neocortex of Fln^KO^ mice, and periventricular heterotopia (PH). Shown are violin plots with individual data point, median, and quartiles. *** p <0.001 by Student’s t-test. Bars: 100 um or as indicated.

To validate that Fln^KO^ neurons in periventricular nodules were less active than those in upper cortical layers, we examined c-Fos, an immediate early gene of the AP-1 transcriptional activator that is rapidly and transiently upregulated in response to neuronal activities(Bullitt, 1990). We found although the overall abundance of c-Fos protein is higher in the cortical tissue of Fln^KO^ than that of control mice (Figure 6C-E), the c-Fos immunoreactivity was presented predominantly by upper cortical neurons, it was notably weaker or undetectable in periventricular neurons (Figure 6C). Furthermore, the low c-Fos in periventricular neurons was accompanied by reduced Camk2, the calcium/calmodulin-dependent protein kinase that is abundantly expressed in excitatory neurons, activated by the NMDA-receptor-mediated calcium elevation, and plays essential roles for synaptic plasticity. Various Camk2 isoforms, including Camk2a, Camk2b, Camk2g, and Camk2n, were among DEGs upregulated in both V-SVZ and upper cortical ROIs of Fln^KO^ compared to control mice (Figure S3, Figure S5), but we found Camk2’s immuno-signals were considerably weaker in periventricular neurons than in upper cortical neurons of the same cortical section (Figure 6F). As the induction of c-Fos can be triggered by neuronal activity-mediated calcium influx and Camk2 activation, the jointed deficiency in c-Fos and Camk2 strongly supports the conclusion of spatial transcriptomics analysis, indicating that in Fln^KO^ mice, periventricular neurons produced in the postnatal V-SVZ are relatively dormant compared to neocortical neurons generated during embryonic development.

An additional feature of periventricular heterotopia nodules is that they contain a lower density of parvalbumin-expressing (PV^+^) GABAergic neurons. These PV^+^ inhibitory interneurons are derived from the ganglionic eminence in embryogenesis and migrate tangentially to the neocortex. In mice, cortical PV^+^ neurons reach a steady state at the end of second postnatal week, adopt their electrophysiological properties at weaning age, and play important roles in regulating the activity of cortical glutamatergic excitatory neurons(Miyamae et al., 2017; Rupert and Shea, 2022). Although the abundance of PV^+^ neurons was higher in the neocortex of Fln^KO^ brains than that of control brains, it was significantly reduced in periventricular nodules (Figure 6G, H). PV^+^ interneurons are important for excitation-inhibition balance, their maturation and coupling to glutamatergic neurons contribute to the development of gamma oscillation and enhance cortical circuit performance(Sohal et al., 2009; Tremblay et al., 2016). Therefore, the enrichment of PV neurons in the neocortex and deficiency of PV^+^ neurons in periventricular heterotopic nodules suggest that neural network activities in Fln^KO^ brains are predominantly conducted by neocortical neurons, whereas the contribution of periventricular neurons born in postnatal and adult V-SVZ is likely minimal.

Despite a molecular function of filamin in cross-linking actin filaments, neither cortical nor periventricular neurons in Fln^KO^ mice showed gross structural aberrations by Golgi-Cox stain and morphology analysis, though neurons in periventricular heterotopia failed to develop proper spatial organization (Figure 6I, Figure S6B). Nonetheless, both heterotopic and cortical neurons in Fln^KO^ mice showed significant increases in dendritic spine density compared to that of cortical neurons in control mice (Figure 6I, J). In addition, the spine morphology of filamin deficient neurons were notably distorted, showing a fragile structure with thinner neck and claw-like glomerular endings. The higher spine density provides a larger number of contact sites for potential synaptic connectivity, and the reduced spine structural strength permit more dynamic synaptic remodeling. These morphological features of dendritic spines are consistent with the overall underdeveloped state of Fln^KO^ neurons with reduced stability but enhanced plasticity and activity.

## Discussion

Although generation of new neurons in the adult brain holds a promise for replenishing neurons lost due to brain injury or degeneration, studies in the past decades have only shown the acquisition of a handful of molecular markers for active NSPCs and neuroblasts or the production of a small number of neurons transiently in the adult brain. Direct evidence for the long-term survival and physiological function of these adult-born neurons remains lacking. As a result of these limitations, whether new neurons can be generated in the adult human brain has been controversial (Duque *et al*., 2022; Nano and Bhaduri, 2022; Paredes et al., 2018). The result of this study demonstrates that a large quantity of neurons resembling neurons of the neocortex can be generated by the postnatal and adult V-SVZ of Fln^KO^ mice, providing a direct evidence on the neurogenic capacity of adult V-SVZ. Given the close genetic and phenotypic resemblance between Fln^KO^ mice and the corresponding human PH condition, our result serves as a proof of concept for further exploring the potential of inducing V-SVZ neurogenesis in the adult human brain.

Despite extensive studies of neurogenesis in the brain, mechanisms underlying the intrinsic and extrinsic control of NSPCs’ neurogenic propensity remain largely elusive. Nonetheless, at the systems-level, it is conceivable that both intrinsic and extrinsic factors driving neuronal differentiation are well synchronized with the developmental state and physiological activities of the brain. This notion is well supported by results of this study. First, our data suggest that NSPCs’ neurogenic propensity is highly regulated by the brain’s metabolic pattern. In fetal development, brain metabolism is predominately mediated by anaerobic glycolysis(Goyal et al., 2014). After birth, the brain starts to receive oxygen-rich blood and gradually switches to aerobic catabolism of glucose. By adult age, brain energetics fully relies on OXPHOS, consuming 25% of glucose intake and 20% of inhaled oxygen in order to maintain neural activities. To meet this energy demand of the adult brain, upregulating genes participating OXPHOS must be necessary for activating the neuronal differentiation of NSPCs as we observed in adult Fln^KO^ mice. In close agreement, previous reports also demonstrated that the transition of neural stem cells to NPCs of the neuronal lineage is accompanied by increased mitochondrial biogenesis(Beckervordersandforth et al., 2017; Zheng et al., 2016). Moreover, our data indicated that generation of periventricular neurons in the V-SVZ of Fln^KO^ mice is correlated with increased expression of a myriad of genes involving neuronal activity, plasticity, and metabolic turnovers in the neocortex. These data imply that in addition to metabolic patterns, neurogenic propensity of NSPCs in the postnatal and adult V-SVZ is well-coordinated with neocortical neurons’ differential state, maturity, and physiological activity. Besides long-distance axonal communications, vitalized V-SVZ and upper cortical layers in Fln^KO^ mice may be functionally connected by vascular circulation. In this case, increased vascularization not only boosts oxidative metabolism and ATP synthesis within V-SVZ but also enhances brain-wide metabolic turnover and function. Therefore, cerebral blood vessels are both an essential neurogenic niche and a prerequisite for neuronal function in the adult brain.

While the increased vascularization acts as the external cue to enhance the energetics and activities of cortical NPCs and neurons, filamin LOF is the intrinsic factor that renders NPCs and neurons to an under-differentiated immature state with high susceptibility and sensitivity to external cues. This is consistent with our earlier finding that filamin LOF results in hyperactive responses to Igf and Vegf due to deficiencies in signaling attenuation in the embryonic V-SVZ(Houlihan *et al*., 2016). Such deficiency suggests that filamin acts as the gatekeeper of cellular and tissue homeostasis. Although filamin has been regarded as an actin stabilizer, the diverse phenotypes of filamin LOF in humans and mice are not simply explained by deficiencies in actin crosslinking. Besides actin, filamin interacts with numerous molecules for cell sensing, signaling, gene expression, and morphogenesis. This allows it to coordinate intricate cell-cell interactions, signaling networks, intracellular structures, and metabolic processes in various tissue environments. This characteristic of filamin is essential for maintaining the homeostasis and stability of cells bombarded by diverse extracellular cues. In line with this, *FLNA* mutations have been shown to affect the cardiovascular, skeletal, lungs, gastro, and connective tissues(Chen and Walsh, 1993; Kyndt *et al*., 2007; Reinstein *et al*., 2013; Robertson, 1993; Sasaki et al., 2019). Recent studies also suggest a dual function of FLNa in cancer promotion and tumor suppression(Savoy and Ghosh, 2013). These are all fully in line with data presented in this study, which demonstrate that filamin loss of function results in the reprograming of a wide range of molecular networks for cell signaling, cell mechanics, gene expression, and metabolism. These changes may weaken the homeostatic stability, resulting in sensitization and activation of NPCs, glia, and neurons at a brain systems-level.

Although epilepsy is the main neurological symptom of *FLNA* mutations, the condition can be asymptomatic and go silently without being detected by magnetic resonance imaging. We have not observed spontaneous seizures in Fln^KO^ mice. However, our spatial transcriptomics data revealed that genes mediating excitatory activities are significantly increased in neurons of upper layers than those of periventricular nodules. This suggests that seizure activities in the human PH condition originate, at least in part, from hyperexcitabilities of neocortical neurons with *FLNA* deficiency. In general, seizure activities often result from shifts in inhibitory and excitatory imbalance towards excessive excitability. In Fln^KO^ mice, the density of inhibitory PV+ neurons was significantly higher in the cortex but significantly lower in periventricular heterotopia. It remains possible that the deficiency of PV+ neurons in periventricular nodules accounts for the hyperexcitability of seizure induction. However, reduced c-Fos and Camk2 in periventricular neurons, along with a transcriptome profile showing decreased neural activity, altogether suggest that heterotopic neurons are unfavorable candidates for hyperactivities. Given that the survival, maturation, and activity of PV+ neurons are influenced substantially by the activity of excitatory circuits, a more plausible interpretation is that the increase in PV+ neurons in the neocortex of Fln^KO^ mice is a compensatory response to a cortical excitation and inhibition imbalance, whereas the low PV+ neurons in the periventricular nodules may be a sign of function immaturity of periventricular excitatory neurons generated by the postnatal and adult V-SVZ.

To date, our understanding of the neurogenic potential and functional significance of the adult V-SVZ is still at a very early stage. There is a pressing need for developing representative in vivo models to uncover the mechanisms for regulating V-SVZ activities. Although heterotopic neurons generated in the V-SVZ of Fln^KO^ mice appear unable to integrate into the cortical circuity and fully acquire activity and function, this mouse model of human condition raises a possibility of inducing V-SVZ neurogenesis by targeting FLNA-mediated NSPC-vascular homeostasis and brain energetics.

### Limitation of the study

Although this study provides direct evidence that the adult V-SVZ retains the ability of generating neurons that resemble those in the neocortex with respect to transcriptome profiles, it is challenging to fully define the underlying mechanism that activates NSPCs in the adult SVZ, since our results suggest it involves coordinated changes in a large number of molecules, cell types, and pathways at the brain systems-level. Likewise, the function of filamin is also multifaceted and complex, which makes it difficult to delineate the precise molecular mechanism underlying functional alterations in V-SVZ and neocortex of Fln^KO^ mice. Our previous developmental study showed that filamin deficient radial glial cells in the VZ undergo sustained EMT, altering the microenvironment of neuronal fate restricted intermediate NPCs in the embryonic SVZ. Although Snail1, the prominent EMT inducer, is significantly upregulated in the V-SVZ of adult Fln^KO^ mice, our transcriptomic analysis did not reveal enrichment of the EMT pathway. Given the numerous molecular interactions of filamin, future studies are needed to delineate the common and differential mechanisms of filamin in NSPCs of the embryonic and adult V-SVZ, neurons of the neocortex, and the neuro-vascular interface that synchronizes neural activities with vascular circulation.

## Methods

### Key resources table

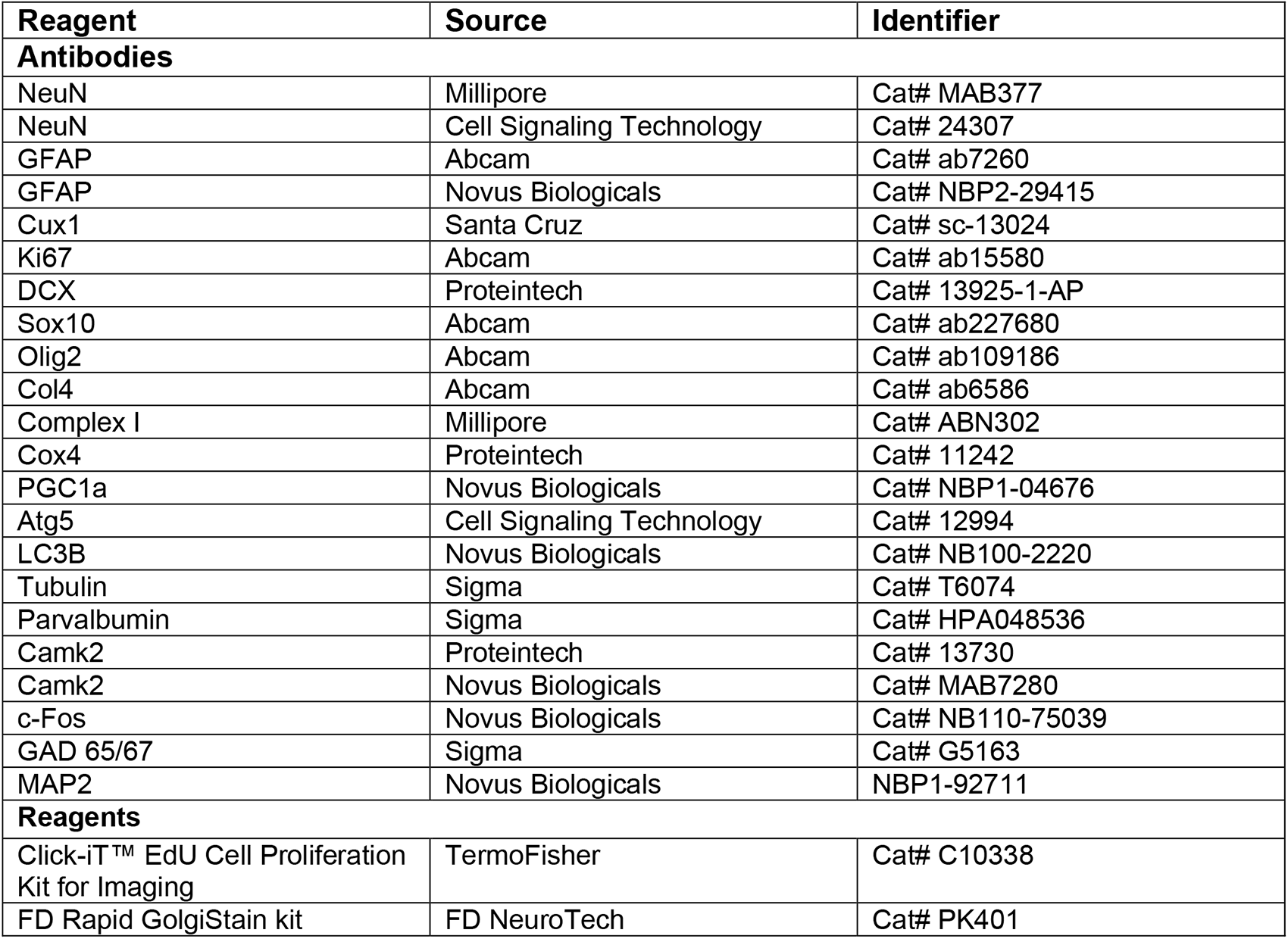

### Resource availability

#### Lead contact

Further information and requests for resources and reagents should be directed to and will be fulfilled by the Lead Contact, Yuanyi Feng.

### Material availability

All the materials generated in this study are available upon reasonable request to the lead contact.

### Mice

Flna floxed (Flna^flox/flox^ female or Flna^flox/y^ male) mice, and Flnb knockout mice (Flnb^-/-^) were generated by conventional mouse embryonic stem cell-based gene targeting (Feng *et al*., 2006; Lu et al., 2007). NSPC-specific Flna conditional knockout mice were generated by crossing Flna^flox/flox^ mice with Tg(Nes-Cre)1Wmz Cre mice(Petersen et al., 2004). The Flna;Flnb compound mutant mice were generated by standard genetic cross. All mice used for this study were housed and bred according to the guidelines approved by the IACUC committees of Uniformed Services University of Health Services in compliance with the AAALAC’s guidelines. Experiments were performed using littermates or age and genetic background matched control and mutant groups in both sexes.

### Fluorescence immunohistological and immunochemical analyses

For immunofluorescence staining of mouse cortical tissue, mouse brains were fixed by transcardial perfusion with PBS and 4% paraformaldehyde and then processed in 12um cryosections or 5 um paraffin sections. After treating with antigen unmasking solutions (Vector Labs), brain sections were blocked with 5% goat serum and incubated with primary antibodies in PBS, 0.05% Triton X100, and 5% goat serum at 4°C overnight, and followed by staining with fluorescence conjugated antibodies and Hoechst 33342. Epifluorescence images were acquired with a Leica CTR 5500 fluorescence, DIC, and phase contrast microscope equipped with the Leica DFC7000T digital camera. Images were imported to Adobe Photoshop and adjusted for brightness and black values.

### GeoMx Digital Spatial Profiling of the Mouse Whole-transcriptome

Preparation of samples for full transcriptome spatial RNA-seq analysis was carried out following the GeoMx DSP slide preparation user manual (MAN-10150). Briefly, brains of three Fln^KO^ and three control mice at 3 months of age were fixed by transcardial perfusion with nuclease-free PBS and 4% paraformaldehyde. From the fixed brains cryosections of 7-μm thickness were placed on a microscope slide. The slide was incubated with mouse whole transcriptome oligonucleotide probe mix overnight, then washed and stained with Alexa Fluo 647 conjugated antibody to NeuN (Cell Signaling Technology), Alexa Fluo 532 conjugated antibody to Gfap, (Novus Biologicals), and DNA dye Syto13. Tissue sections were then loaded into the GeoMx DSP platform by which immunofluorescence signals were visualized by three colors (Green = Gfap; Red = NeuN; blue = Syto13) for selecting Regions-Of-Interests (ROIs). Circular geometric ROIs of ∼100um in diameter were selected from V-SVZ or upper cortical layers of control and Fln^KO^ brains, as well as periventricular nodules of Fln^KO^ brains. Oligonucleotide probes in each ROI were released by UV light and collected in separate wells of a microtiter plate. Photo-released nucleotides were used to generate sequencing libraries by PCR, during which Illumina i5 and i7 dual-indexing primers were added to uniquely index each ROI. PCR reactions were purified by AMPure beads twice. Library concentration was determined by a Qubit fluorometer and libraries were paired-end sequenced on a NextSeq 500 system (Illumina). The sequencing results were processed through standard GeoMx NGS Pipeline according to GeoMx DSP NGS Readout User Manual (MAN-10153-01), during which raw sequencing FASTQ files were converted to digital count conversion (DCC) files and further processed to obtain data for each target probe in each ROI by subtracting the mean of background and normalizing to Q3 of all targets. The ROIs were categorized according to genotype and spatial groups. Differential gene expression analyses were performed on different groups of ROIs by Student’s t test with Benjamini-Hochberg adjustment. Adjusted p values of < 0.05 were regarded significant.

### Ultra-Structural Analysis

Mouse brains were fixed with EM grade formaldehyde (2%) and glutaraldehyde (2%) in PBS by transcardial perfusion. The neocortex and V-SVZ of perfused brains were dissected, cut to pieces of < 1 mm^3^ in size, immersed in the same fixative overnight, and washed three times in PBS. Samples were then washed 3x 15min in cacoldylate buffer (CB, 0.1M, pH 7.4) to remove phosphate ions and subsequently immersed in 2% OsO_4_ in CB for 1 hour. Following 3x15 min washes in CB, samples were dehydrated in a graduated series of ethanol, infiltrated with Spurr’s epoxy resin (Electron Microscopy Sciences, Hatfield, PA), and polymerized at 70℃ for 10 hours. Thin sections (70-90nm) were cut on a Leica Ultracut UC-6 ultramicrotome (Leica Microsystems, Wetzlar, Germany). Sections were collected on 3mm copper grids and subsequently stained for 30min with 2% aqueous uranyl acetate and for 5min with Reynold’s lead citrate. The stained brain specimens were examined with a JEOL JEM-1011 transmission electron microscope. Images were acquired by an Advanced Microscopy Techniques 4MP digital camera (AMT Corp) and saved as TIFF files. Mitochondria quantification was performed by manually tracing the outer mitochondrial membrane using ImageJ.

### Golgi-Cox Staining for Neuronal Morphology and Dendritic Spine Analysis

Mice were euthanized with CO_2_; brains were quickly dissected, rinsed with deionized water, immersed in impregnation solution, and processed using FD Rapid GolgiStain kit (FD NeuroTechnologies) according to manufacturer’s instructions. Stained sections were examined under a Leica DM5000 light microscope. Pyramidal neurons in the cerebral cortex and hippocampus regions were imaged with a 40x objective and photographed. For dendritic spine density analysis, 16-20 pyramidal neurons in neocortical layer II/III of each mouse were randomly selected for assessment. The number of spines per 10 micrometers in secondary apical dendrites (branched from primary dendrites arising from the soma) was scored using the NIH Image J software.

### Immunoblotting

Immunoblotting of total cell or tissue proteins was performed by extracting with boiling 2x SDS PAGE sample buffer (62.5 mM Tris-HCl, pH 6.8, 2.5% SDS, 0.7135 M β-mercaptoethanol, 10% glycerol, 0.002% Bromophenol Blue) to fully dissolve the tissue proteins, heating at 95°C for 10 min to ensure protein denaturation, and passing through syringes with a 29-gauge needle three times to sheer nuclear DNA and obtain homogenous extracts. 10-30 ug of total proteins were used for each immunoblotting analysis. The loadings were adjusted and normalized by the total protein content according to Coomassie blue stain of the gel after SDS PAGE and by the level of housekeeping proteins.

### Quantification and statistical analysis

No statistical methods were used to predetermine sample size, while all experiments were performed with a minimum of three biological replicates and all cell counts were obtained from at least ten random fields. The experiments were not randomized; the investigators were not blinded to the sample allocation and data acquisition during experiments but were blinded in performing quantitative analyses of immunohistological images using the NIH ImageJ software.

All statistical analyses were done using GraphPad Prism 9.0 software. Data were analyzed by one-way ANOVA or unpaired two-tailed Student’s t tests for comparing differences between different genotypes. Differences were considered significant with a p value < 0.05. Data distribution was assumed to be normal with the exception of survival analysis, but this was not formally tested.

## Data availability

All raw and processed data used in this article will be available upon request in writing to the corresponding author.

## Author contributions

Y.F. conceptualized the project, designed and performed the experiments, interpreted the results, and wrote the manuscript. C.A.K. performed experiments and data analysis; C.C.L. performed the experiments and data analysis; D.P.M. assisted with ultrastructural imaging analyses, and X.Z. assisted with experiments.

## Disclaimer

The opinions, interpretations, conclusions and recommendation are those of the authors and are not necessarily endorsed by the U.S. Army, Department of Defense, the U.S. Government or the Uniformed Services University of the Health Sciences. The use of trade names does not constitute an official endorsement or approval of the use of reagents or commercial hardware or software. This document may not be cited for purposes of advertisement.

