## Supplemental Figures for "Sustained Generation of Neurons Destined for Neocortex with Oxidative Metabolic Upregulation upon Filamin Abrogation"

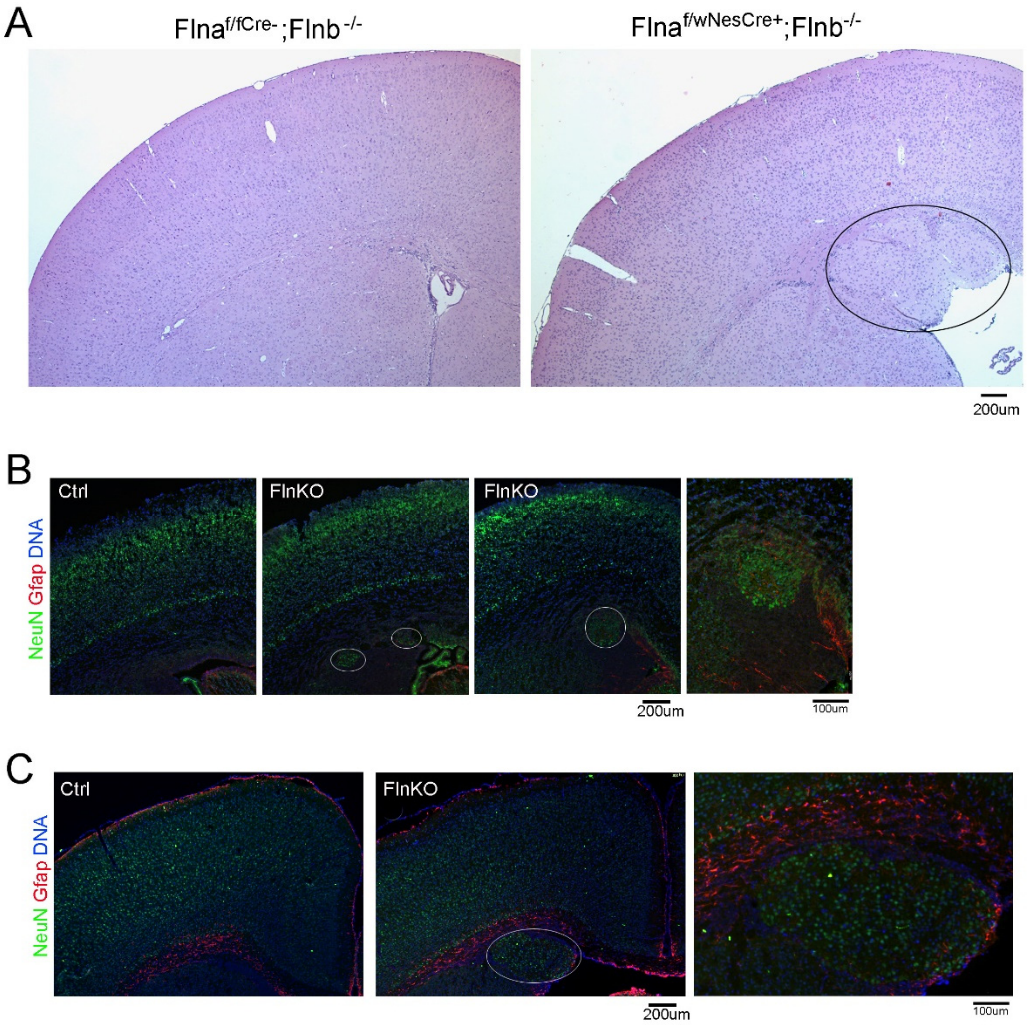


**Figure S1. Periventricular Heterotopia and their postnatal growth in $Fln^{KO}$ mice.**

A. H&E stained brain sections of a Flnb homozygous knockout mice and a female heterozygous Flna cKO compounded with Flnb homozygous KO mice at 2 months of age. The position of circles indicates PH

B. Representative images of NeuN (green) and Gfap (red) double immunostained brain sections of $Fln^{KO}$ or control mice shortly after birth (P0). Nuclear DNA was stained with Hoechst 33342 and shown in blue. The position of circles indicates PH.

C. Representative images of NeuN (green) and Gfap (red) double immunostained brain sections of $Fln^{KO}$ or control mice at P7. Nuclear DNA was stained with Hoechst 33342 and shown in blue. The position of circles indicates PH.

Bars: 100um or 200um as indicated.

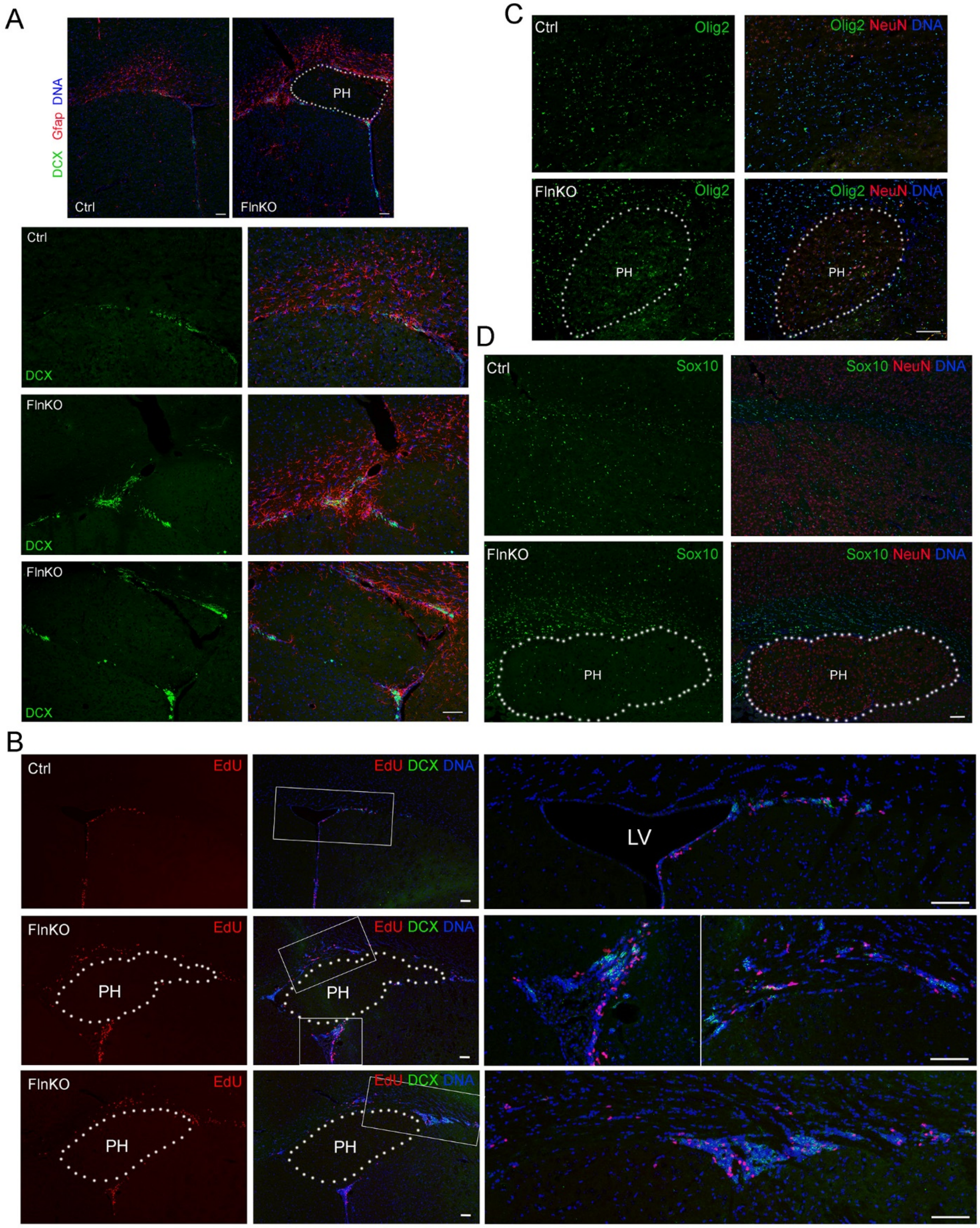

A
DCX Gfap DNA
PH
Ctrl
FlnKO
Ctrl
DCX
FlnKO
DCX
FlnKO
DCX
B
Ctrl
EdU
EdU DCX DNA
LV
FlnKO
EdU
PH
EdU DCX DNA
PH
FlnKO
EdU
PH
EdU DCX DNA
PH
C
Ctrl
Olig2
Olig2 NeuN DNA
FlnKO
Olig2
PH
Olig2 NeuN DNA
PH
D
Ctrl
Sox10
Sox10 NeuN DNA
FlnKO
Sox10
PH
Sox10 NeuN DNA
PH

**Figure S2. Increased cellular activities in the V-SVZ of $Fln^{KO}$ mice**

A. Representative images of DCX (green) and Gfap (red) double immunostained brain sections of $Fln^{KO}$ or control mice at 3 months of age. Nuclear DNA was stained with Hoechst 33342 and shown in blue. The position of PH is marked by dotted lines.

B. Representative images of brain sections stained by EdU (red) and immunostained by anti-DCX (green) of $Fln^{KO}$ or control mice at 3 months of age. Nuclear DNA was stained with Hoechst 33342 and shown in blue. The position of PH is marked by dotted lines.

C. Representative images of Olig2 (green) and NeuN (red) double immunostained brain sections of $Fln^{KO}$ or control mice at 3 months of age. Nuclear DNA was stained with Hoechst 33342 and shown in blue. The position of PH is marked by dotted lines.

D. Representative images of Sox10 (green) and NeuN (red) double immunostained brain sections of $Fln^{KO}$ or control mice at 2 months of age. Nuclear DNA was stained with Hoechst 33342 and shown in blue. The position of PH is marked by dotted lines.

Bars: 100 um.

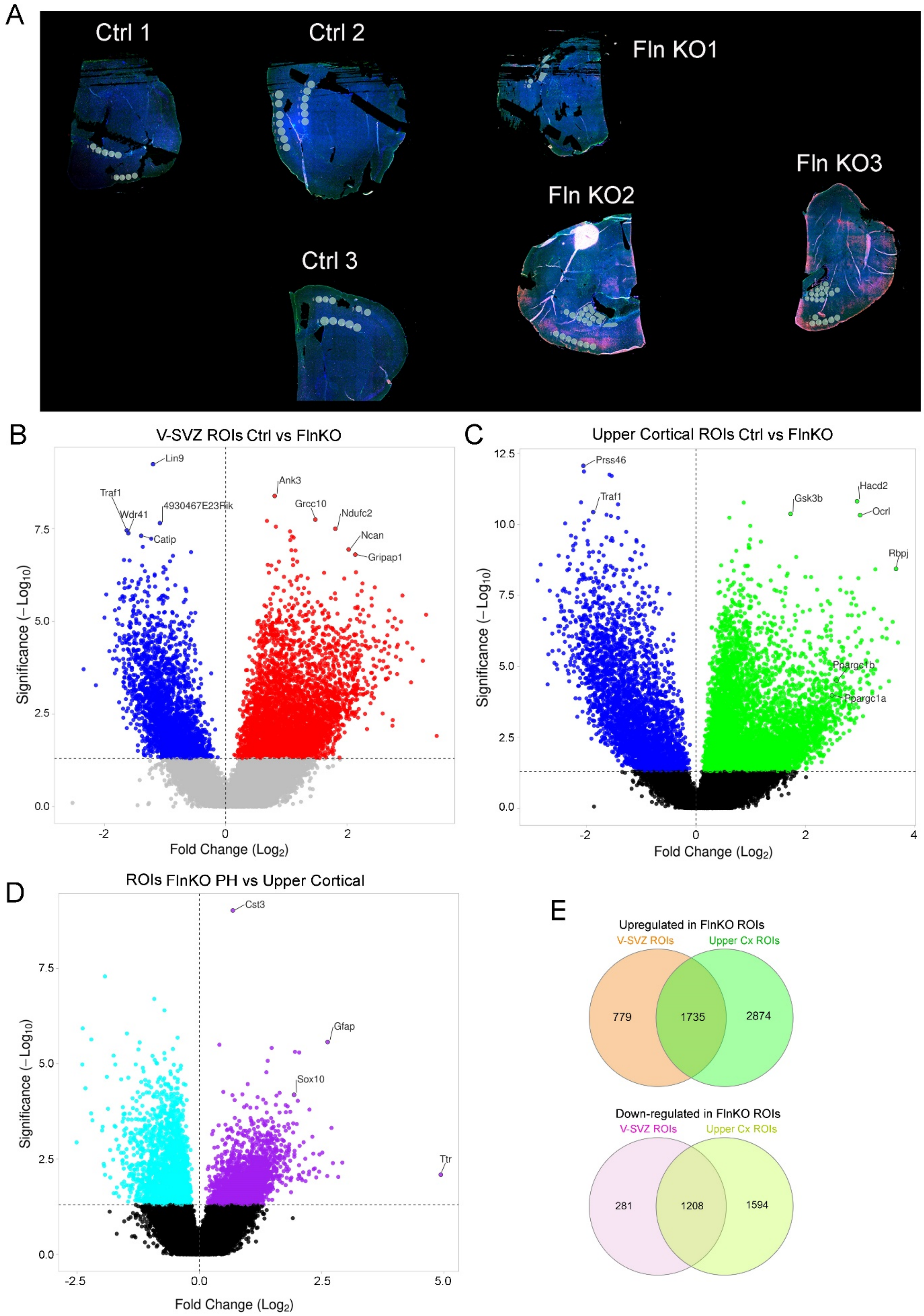

A
Ctrl 1
Ctrl 2
Fln KO1
Fln KO3
Fln KO2
Ctrl 3
B
V-SVZ ROIs Ctrl vs FlnKO
Lin9
Traf1
Wdr41
4930467E23Rik
Catip
Ank3
Grcc10
Ndufc2
Ncan
Gripap1
Significance (−Log10)
Fold Change (Log2)
C
Upper Cortical ROIs Ctrl vs FlnKO
Prss46
Traf1
Gsk3b
Hacd2
Ocrl
Rbpj
Ppargc1b
Ppargc1a
Significance (−Log10)
Fold Change (Log2)
D
ROIs FlnKO PH vs Upper Cortical
Cst3
Gfap
Sox10
Ttr
Significance (−Log10)
Fold Change (Log2)
E
Upregulated in FlnKO ROIs
V-SVZ ROIs
Upper Cx ROIs
779
1735
2874
Down-regulated in FlnKO ROIs
V-SVZ ROIs
Upper Cx ROIs
281
1208
1594

**Figure S3. Spatial mouse whole-transcriptome analyses by GeoMx Digital Spatial Profiling**

A. Image of the slide analyzed by GeoMx DSP. The slide contains brain sections from three control mice and three $Fln^{KO}$ mice at 3-4 months of age. Shown are locations and distribution of the 76 regions of interests (ROIs) being analyzed. These include 16 ROIs in the V-SVZ of control mice, 15 ROIs in the V-SVZ of $Fln^{KO}$ mice, 16 ROIs in the upper cortical layers of control mice, 15 ROIs in the upper cortical layers of $Fln^{KO}$ mice, and 14 ROIs in periventricular heterotopia nodules of $Fln^{KO}$ mice.

B. Volcano plot of DEGs between ROIs in V-SVZ of $Fln^{KO}$ and control mice.

C. Volcano plot of DEGs between ROIs in upper cortical layers of $Fln^{KO}$ and control mice.

D. Volcano plot of DEGs between ROIs in upper cortical layers and periventricular nodules of $Fln^{KO}$ mice.

E. Venn diagram of DEGs upregulated or down-regulated in ROIs in V-SVZ and upper cortical layers of $Fln^{KO}$ mice relative to control mice.

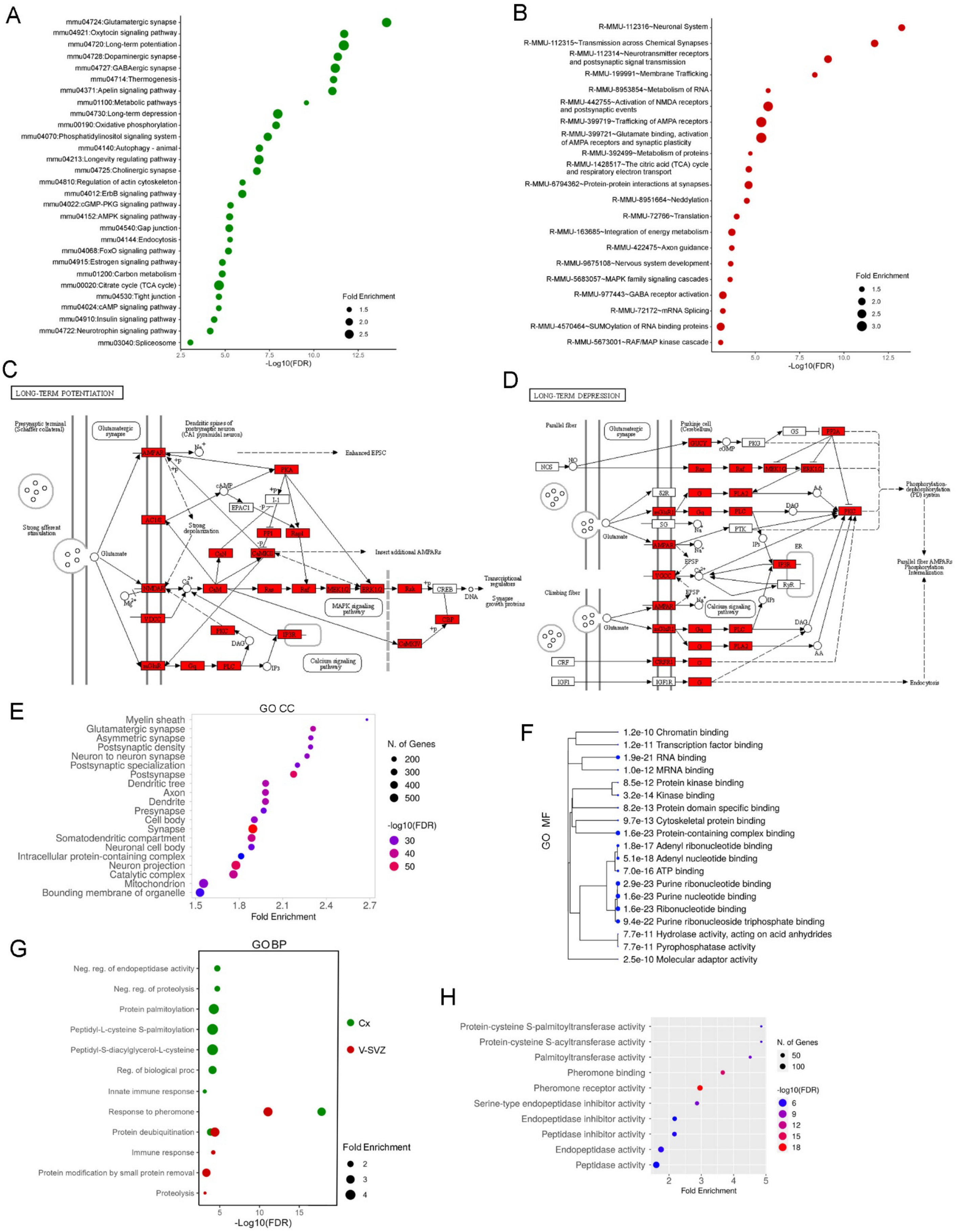
A
mmu04724:Glutamatergic synapse
mmu04921:Oxytocin signaling pathway
mmu04720:Long-term potentiation
mmu04728:Dopaminergic synapse
mmu04727:GABAergic synapse
mmu04714:Thermogenesis
mmu04371:Apelin signaling pathway
mmu01100:Metabolic pathways
mmu04730:Long-term depression
mmu00190:Oxidative phosphorylation
mmu04070:Phosphatidylinositol signaling system
mmu04140:Autophagy - animal
mmu04213:Longevity regulating pathway
mmu04725:Cholinergic synapse
mmu04810:Regulation of actin cytoskeleton
mmu04012:ErbB signaling pathway
mmu04022:cGMP-PKG signaling pathway
mmu04152:AMPK signaling pathway
mmu04540:Gap junction
mmu04144:Endocytosis
mmu04068:FoxO signaling pathway
mmu04915:Estrogen signaling pathway
mmu01200:Carbon metabolism
mmu00020:Citrate cycle (TCA cycle)
mmu04530:Tight junction
mmu04024:cAMP signaling pathway
mmu04910:Insulin signaling pathway
mmu04722:Neurotrophin signaling pathway
mmu03040:Spliceosome
Fold Enrichment
1.5
2.0
2.5
2.5
5.0
7.5
10.0
12.5
-Log10(FDR)
B
R-MMU-112316~Neuronal System
R-MMU-112315~Transmission across Chemical Synapses
R-MMU-112314~Neurotransmitter receptors and postsynaptic signal transmission
R-MMU-199991~Membrane Trafficking
R-MMU-8953854~Metabolism of RNA
R-MMU-442755~Activation of NMDA receptors and postsynaptic events
R-MMU-399719~Trafficking of AMPA receptors
R-MMU-399721~Glutamate binding, activation of AMPA receptors and synaptic plasticity
R-MMU-392499~Metabolism of proteins
R-MMU-1428517~The citric acid (TCA) cycle and respiratory electron transport
R-MMU-6794362~Protein-protein interactions at synapses
R-MMU-8951664~Neddylation
R-MMU-72766~Translation
R-MMU-163685~Integration of energy metabolism
R-MMU-422475~Axon guidance
R-MMU-9675108~Nervous system development
R-MMU-5683057~MAPK family signaling cascades
R-MMU-977443~GABA receptor activation
R-MMU-72172~mRNA Splicing
R-MMU-4570464~SUMOylation of RNA binding proteins
R-MMU-5673001~RAF/MAP kinase cascade
Fold Enrichment
1.5
2.0
2.5
3.0
5.0
7.5
10.0
12.5
-Log10(FDR)
C
LONG-TERM POTENTIATION
Presynaptic terminal (Schaffer collateral)
Glutamatergic synapse
Dendritic spines of postsynaptic neuron (CA1 pyramidal neuron)
Strong afferent stimulation
Glutamate
AMPAR
NMDAR
VDCC
mGluR
Gq
PLC
IP3
DAG
PKC
IP3R
CaM
CaN
CaMKII
AC1/8
cAMP
EPAC1
PKA
I-1
PP1
Rap1
Strong depolarization
Enhanced EPSC
Insert additional AMPARs
Ras
Raf
MEK1/2
ERK1/2
Rsk
CREB
CBP
CaMKIV
DNA
Transcriptional regulators
Synapse growth proteins
MAPK signaling pathway
Calcium signaling pathway
D
LONG-TERM DEPRESSION
Parallel fiber
Glutamatergic synapse
Purkinje cell (Cerebellum)
NOS
NO
GUCY
cGMP
PKG
GS
PP2A
Ras
Raf
MEK1/2
ERK1/2
S2R
mGluR1
SG
AMPAR
Gq
PLC
PLA2
PTK
AA
DAG
IP3
IP3R
RyR
ER
VGCC
EPSP
Calcium signaling pathway
PKC
Phosphorylation-dephosphorylation (PD) system
Parallel fiber AMPARs Phosphorylation Internalization
Climbing fiber
Glutamate
CRF
CRFR1
IGF1
IGF1R
Endocytosis
E
GO CC
Myelin sheath
Glutamatergic synapse
Asymmetric synapse
Postsynaptic density
Neuron to neuron synapse
Postsynaptic specialization
Postsynapse
Dendritic tree
Axon
Dendrite
Presynapse
Cell body
Synapse
Somatodendritic compartment
Neuronal cell body
Intracellular protein-containing complex
Neuron projection
Catalytic complex
Mitochondrion
Bounding membrane of organelle
1.5
1.8
2.1
2.4
2.7
Fold Enrichment
N. of Genes
200
300
400
500
-log10(FDR)
30
40
50
F
GO MF
1.2e-10 Chromatin binding
1.2e-11 Transcription factor binding
1.9e-21 RNA binding
1.0e-12 MRNA binding
8.5e-12 Protein kinase binding
3.2e-14 Kinase binding
8.2e-13 Protein domain specific binding
9.7e-13 Cytoskeletal protein binding
1.6e-23 Protein-containing complex binding
1.8e-17 Adenyl ribonucleotide binding
5.1e-18 Adenyl nucleotide binding
7.0e-16 ATP binding
2.9e-23 Purine ribonucleotide binding
1.6e-23 Purine nucleotide binding
1.6e-23 Ribonucleotide binding
9.4e-22 Purine ribonucleoside triphosphate binding
7.7e-11 Hydrolase activity, acting on acid anhydrides
7.7e-11 Pyrophosphatase activity
2.5e-10 Molecular adaptor activity
G
GO BP
Neg. reg. of endopeptidase activity
Neg. reg. of proteolysis
Protein palmitoylation
Peptidyl-L-cysteine S-palmitoylation
Peptidyl-S-diacylglycerol-L-cysteine
Reg. of biological proc
Innate immune response
Response to pheromone
Protein deubiquitination
Immune response
Protein modification by small protein removal
Proteolysis
5
10
15
-Log10(FDR)
Cx
V-SVZ
Fold Enrichment
2
3
4
H
Protein-cysteine S-palmitoyltransferase activity
Protein-cysteine S-acyltransferase activity
Palmitoyltransferase activity
Pheromone binding
Pheromone receptor activity
Serine-type endopeptidase inhibitor activity
Endopeptidase inhibitor activity
Peptidase inhibitor activity
Endopeptidase activity
Peptidase activity
2
3
4
5
Fold Enrichment
N. of Genes
50
100
-log10(FDR)
6
9
12
15
18

**Figure S5. Functional enrichment analyses of DEGs in V-SVZ or Upper Cortical Layer ROIs between Fln$^{KO}$ and control mice.**

A. KEGG pathway enrichment analysis of upregulated DEGs in upper cortical layer ROIs of Fln$^{KO}$ mice compared to those of control mice.

B. Reactome pathway enrichment analysis of upregulated DEGs in upper cortical layer ROIs of Fln$^{KO}$ mice compared to those of control mice.

C. Schematic representation of KEGG pathway "Long-Term Potentiation". Genes highlighted in red were upregulated across upper cortical layer ROIs of Fln$^{KO}$ mice relative to those of control mice.

D. Schematic representation of KEGG pathway "Long-Term Depression". Genes highlighted in red were upregulated across upper cortical layer ROIs of Fln$^{KO}$ mice relative to those of control mice.

E. Gene Ontology Cellular Component (CC) terms enriched in DEGs upregulated across upper cortical layer ROIs in Fln$^{KO}$ mice compared to those of control mice.

F. Gene Ontology Molecular Function (MF) terms enriched in DEGs upregulated across upper cortical layer ROIs in Fln$^{KO}$ mice compared to those of control mice.

G. Gene Ontology Biological Processes (BP) terms enriched in DEGs downregulated across V-SVZ or upper cortical layer ROIs in Fln$^{KO}$ mice compared to those of control mice.

H. Gene Ontology Molecular Function (MF) terms enriched in DEGs downregulated upper cortical layer ROIs in Fln$^{KO}$ mice compared to those of control mice.

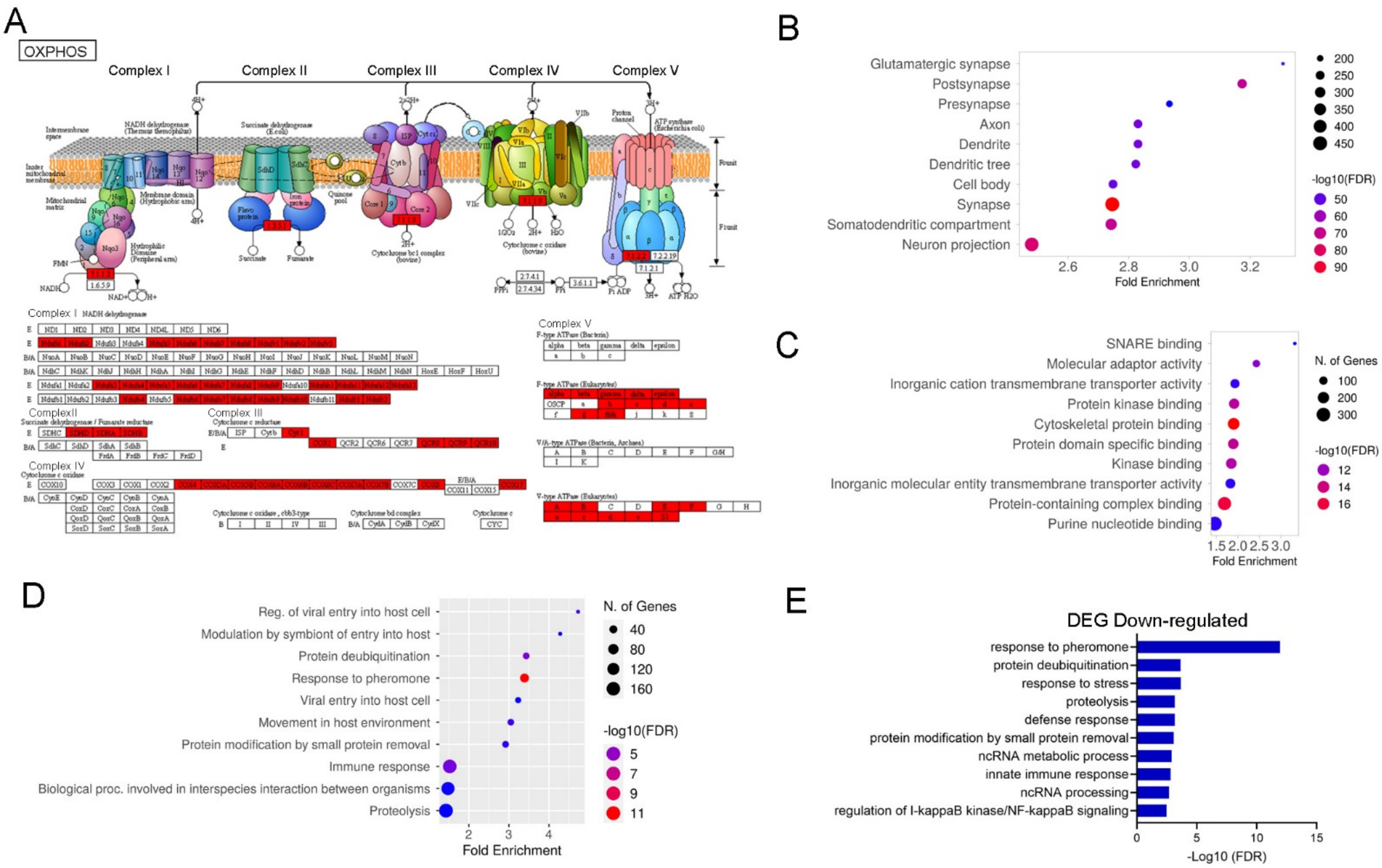


**Figure S4. Functional enrichment analyses of DEGs in V-SVZ ROIs between Fln$^{KO}$ and control mice.**

A. Schematic representation of KEGG pathway "Oxidative Phosphorylation". Genes highlighted in red were upregulated across V-SVZ ROIs of Fln$^{KO}$ mice relative to those of control mice. Some highlighted DEGs were upregulated in both V-SVZ and upper cortical ROIs of Fln$^{KO}$ mice. No genes within the Oxidative Phosphorylation pathway was downregulated in Fln$^{KO}$ mice.

B. Gene Ontology Cellular Component (CC) terms enriched in DEGs upregulated across V-SVZ ROIs in Fln$^{KO}$ mice compared to those of control mice.

C. Gene Ontology Molecular Function (MF) terms enriched in DEGs upregulated across V-SVZ ROIs in Fln$^{KO}$ mice compared to those of control mice.

D, E. Gene Ontology Biological Processes (BP) terms enriched in DEGs down-regulated across V-SVZ ROIs in Fln$^{KO}$ mice compared to those of control mice.

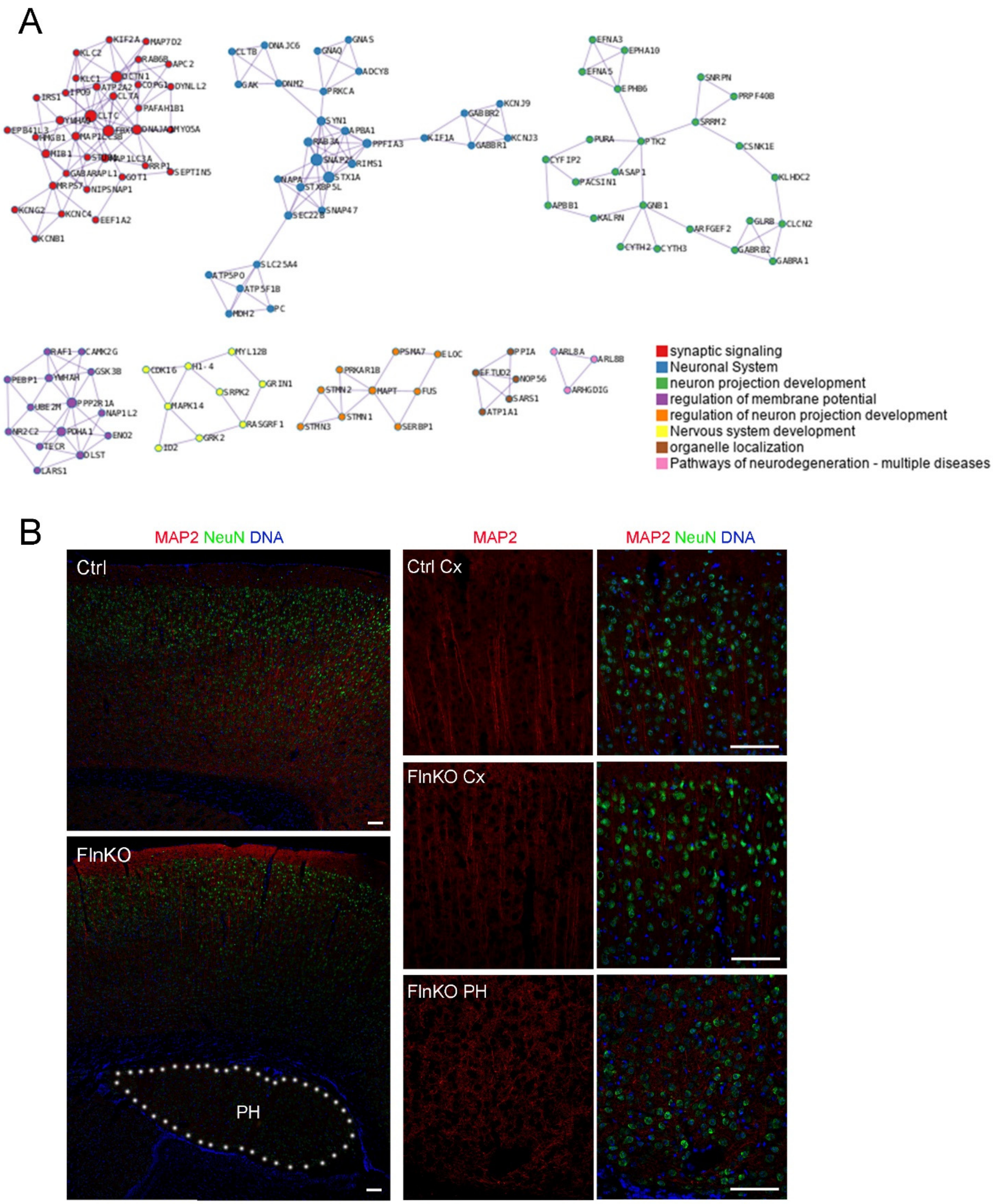
A
synaptic signaling
Neuronal System
neuron projection development
regulation of membrane potential
regulation of neuron projection development
Nervous system development
organelle localization
Pathways of neurodegeneration - multiple diseases
B
MAP2 NeuN DNA
Ctrl
FlnKO
PH
MAP2
MAP2 NeuN DNA
Ctrl Cx
FlnKO Cx
FlnKO PH

**Figure S6. Periventricular neurons show down-regulation in genes for neural activity and lack of polarized dendritic organization.**

A. Network of genes downregulated in ROIs across periventricular nodules compared to ROIs across upper cortical layers of $Fln^{KO}$ mice.

B. Representative images of Map2 (red) and NeuN (green) double immunostained brain sections of $Fln^{KO}$ or control mice at 3 months of age. Nuclear DNA was stained with Hoechst 33342 and shown in blue. The position of PH is marked by dotted lines.
